# Deconvolving organogenesis in space and time via spatial transcriptomics in thick tissues

**DOI:** 10.1101/2024.09.24.614640

**Authors:** Soichiro Asami, Chenshuo Yin, Luis A. Garza, Reza Kalhor

## Abstract

Organ development is guided by a space-time landscape that constraints cell behavior. This landscape is challenging to characterize for the hair follicle – the most abundant mini organ – due to its complex microscopic structure and asynchronous development. We developed 3DEEP, a tissue clearing and spatial transcriptomic strategy for characterizing tissue blocks up to 400 µm in thickness. We captured 371 hair follicles at different stages of organogenesis in 1 mm^3^ of skin of a 12-hour-old mouse with 6 million transcripts from 81 genes. From this single time point, we deconvoluted follicles by age based on whole-organ molecular pseudotimes to animate a stop-motion 3D atlas of follicle development along its trajectory. We defined molecular stages for hair follicle organogenesis and characterized the order of emergence for its structures, differential signaling dynamics at its top and bottom, morphogen shifts preceding and accompanying structural changes, and series of structural changes leading to the formation of its canal and opening. We further found that hair follicle stem cells and their niche are established and stratified early in organogenesis, before the formation of the hair bulb. Overall, this work demonstrates the power of increased depth of spatial transcriptomics to provide a four-dimensional analysis of organogenesis.

## Introduction

The hair follicle is the smallest, most numerous organ in the body and a defining feature of mammals in evolution (Litman and Stein 2023). Its formation, similar to that of many larger organs, involves prototypic coordination between different germ layers (Schneider et al. 2009). It is one of the few human organs capable of near-infinite, life-long regeneration involving cycles of growth, regression, and quiescence (Welle 2023). Its dysregulation can cause medical conditions such as Alopecia (a form of excessive hair loss) and Hirsutism (a form of excessive hair growth), among others (Wolff et al. 2016). Given its abundance and cycling nature, the hair follicle is a unique model of organ development and regeneration in mammals.

Three distinct lineages coordinate to form the diversity of cells and structures in the hair follicle in the latter half of embryogenesis: epidermis, neural crest, and dermis (Saxena et al. 2019). The epidermis, which is ectoderm-derived, contributes the main hair follicle cells known as keratinocytes. Keratinocytes invaginate into the dermis during follicle formation, creating the bulk of the hair follicle’s structure and ultimately generating the hair fiber itself. The neural crest, which is also ectoderm-derived, contributes fibroblasts and melanocytes that migrate into the follicle from the skin surface, forming melanocyte stem cells and mature melanocytes that intercalate multiple keratinocytes-derived layers of the follicle (Vandamme and Berx 2019; Myung et al. 2022). These cells are responsible for the pigmentation of the hair. Finally, the dermis, which is mesoderm-derived, contributes fibroblasts that form the dermal sheath around the follicle and the dermal papilla at the base of its bulb (Martino et al. 2021). These fibroblasts provide a niche for hair follicle formation and are believed to regulate its maintenance and regeneration later in life (Yang et al. 2017; Heitman et al. 2020).

Despite its abundance and accessibility, understanding hair follicle formation and morphogenesis at a cellular level remains challenging for two reasons. First, hair follicles develop asynchronously during the fetal and early postnatal periods. At any point in this window, the skin contains thousands of follicles from a spectrum of stages, making it challenging to reconstruct their developmental trajectory. For example, single-cell RNA sequencing (scRNA-seq), or other approaches that rely on tissue dissociation, have captured the diversity of cell types and cell states within hair follicles during development (Ge et al. 2020; Morita et al. 2021); however, they are unable to identify the cells that originate from follicles in the same stage of development and thus cannot deconvolve their developmental trajectory. This asynchrony, therefore, necessitates using imaging-based approaches, which are hindered by the second challenge. Hair follicle structure is micron-scale and non-repetitive along all three axes. This structure is difficult to analyze in full using conventional histological and imaging methods that rely on capturing 2D planes. 2D methods perform best for organs that are large enough to be serially sectioned (e.g., the heart) for 3D reconstruction or those that are repetitive along at least one axis and can be captured by choosing the 2D plane perpendicular to that axis (e.g., a nerve cord). While state-of-the-art spatial transcriptomic approaches have been effective at profiling many other organs (Lee et al. 2014; Rodriques et al. 2019; Xia et al. 2019; Gyllborg et al. 2020; Kebschull et al. 2020; Liu et al. 2020; Srivatsan et al. 2021; Dardani et al. 2022; Lake et al. 2023; Yao et al. 2023; Kalhor et al. 2024), they are not suitable for the hair follicle as most are effectively limited to thin 2D planes. While there has been progress in extending spatial transcriptomics to thicker tissue sections (Wang et al. 2018; Wang et al. 2021), what remains lacking is the combination of scalability to a large number of targets, transcript capture efficiency, and fluorescence signal-to-noise levels required for characterizing hair follicles that are hundreds of microns deep within the skin.

To address these challenges, we first developed 3DEEP-FISH, a new spatial transcriptomic approach for multiplexed detection of target transcripts in tissue blocks as thick as 400 microns. We applied 3DEEP-FISH to 85 genes related to hair follicle development in newborn mouse skin, detecting 6,601,822 transcripts in 371 full hair follicles at different stages of organogenesis in a cubic millimeter of skin volume at a single timepoint. We found that the molecular composition of each follicle closely reflects its developmental stage and length, allowing us to leverage their asynchronicity as an asset by arranging the 371 follicles along their developmental trajectory, effectively creating a stop-motion animation of hair follicle organogenesis in space and time. This animated atlas unravels the order of emergence and dynamics of expansion for different structures within the hair follicle. It revealed extensive shifts in morphogen activity within hair follicle structures over the course of its organogenesis, identified a steady morphogen signaling center near the top of the follicle, and captured the dynamic changes of the dermal papilla, the morphogen signaling center at the follicle’s base. We further characterized the structural, cellular, and molecular events involved in the formation of the hair follicle opening and canal. Finally, we analyzed the positions of melanocyte and keratinocyte stem cells within the follicle and found that they occupy stratified spatial positions at their eventual niche – known as the bulge – in the earliest stage of organogenesis. Overall, this work establishes a new approach for spatial transcriptomic profiling in thick tissue blocks, provides a spatiotemporal (4D) organogenesis atlas of hair follicles, and establishes fundamental insights into the formation of hair follicles.

## Results

### Enabling spatial transcriptomics in thick tissue blocks by removing genomic DNA

To characterize hair follicle formation, we initially attempted to optimize existing gel embedding and tissue clearing approaches (Chung et al. 2013; Sylwestrak et al. 2016; Alon et al. 2021) and used extended incubation times to perfuse tissue samples with transcript-specific padlock probes and reagents for rolling-circle amplification (RCA) (**Fig. 1A**). Using formalin-fixed 400-micron mouse liver blocks as a test, we first permeabilized and cleared the tissue with SDS solution for five days while concurrently diffusing in a padlock probe targeting the abundant *Apoa2* RNA transcript and a primer for RCA of the probe. The RCA primer was modified on its 5’ end with an acrydite moiety, allowing us to perfuse in acrylamide monomers and crosslink them to create a hydrogel matrix to which the target transcripts and their probes are affixed through the RCA primer (**Fig. 1A**). We then treated the hydrogel-embedded tissue with Proteinase K to remove proteins that can inhibit downstream enzymatic steps. After circularizing the hybridized padlock probes using SplintR ligase, we carried out RCA with Phi29 DNA polymerase to create rolling circle amplicons and detected these amplicons with a fluorescent probe. The results showed that amplicon density decayed rapidly with depth within the tissue to the point that few amplicons were generated beyond 50 microns from the tissue surface (**Fig. 1B,C**). Because the incubation times at each step had been long enough for diffusion deep into the tissue, these results suggested that either a diffusion barrier or a non-specific sink for some of the reagents remained within the tissue. However, the combination of SDS and proteinase K treatment is expected to remove lipids and proteins. We thus hypothesized that genomic DNA limited the penetration of our reagents into the tissue, likely by acting as a non-specific sink for some enzymes. Revising the protocol to include DNase I treatment prior to padlock hybridization to mRNA (**Fig. 1A**), we obtained a dramatic improvement: *Apoa2* amplicons were detected throughout the 400-micron depth of liver blocks with a higher capture efficiency (**Fig. 1B,C**). Moreover, fluorescence signal-to-noise ratio (SNR) was consistently above 7 throughout the depth of the specimen with a 25X objective (**Fig. 1D**). The ability to use medium magnification objectives (i.e., 20–25X) enables scanning large volumes rapidly and accurately. Overall, these results establish a strategy for spatial transcriptomic characterization of thick tissue blocks by combining clearing and hydrogel embedding with genomic DNA removal. We call this approach <u>3D</u> Hydrogel-<u>E</u>mbedding for <u>E</u>xpression <u>P</u>rofiling (3DEEP) with <u>F</u>luorescence <u>I</u>n <u>S</u>itu <u>H</u>ybridization (FISH). 3DEEP-FISH offers a combination of high SNR, capture efficiency, and scalability to a large number of genes in thick tissue specimens that, as we demonstrate below, enables spatiotemporal characterization of hair follicle formation.

**Figure 1.**
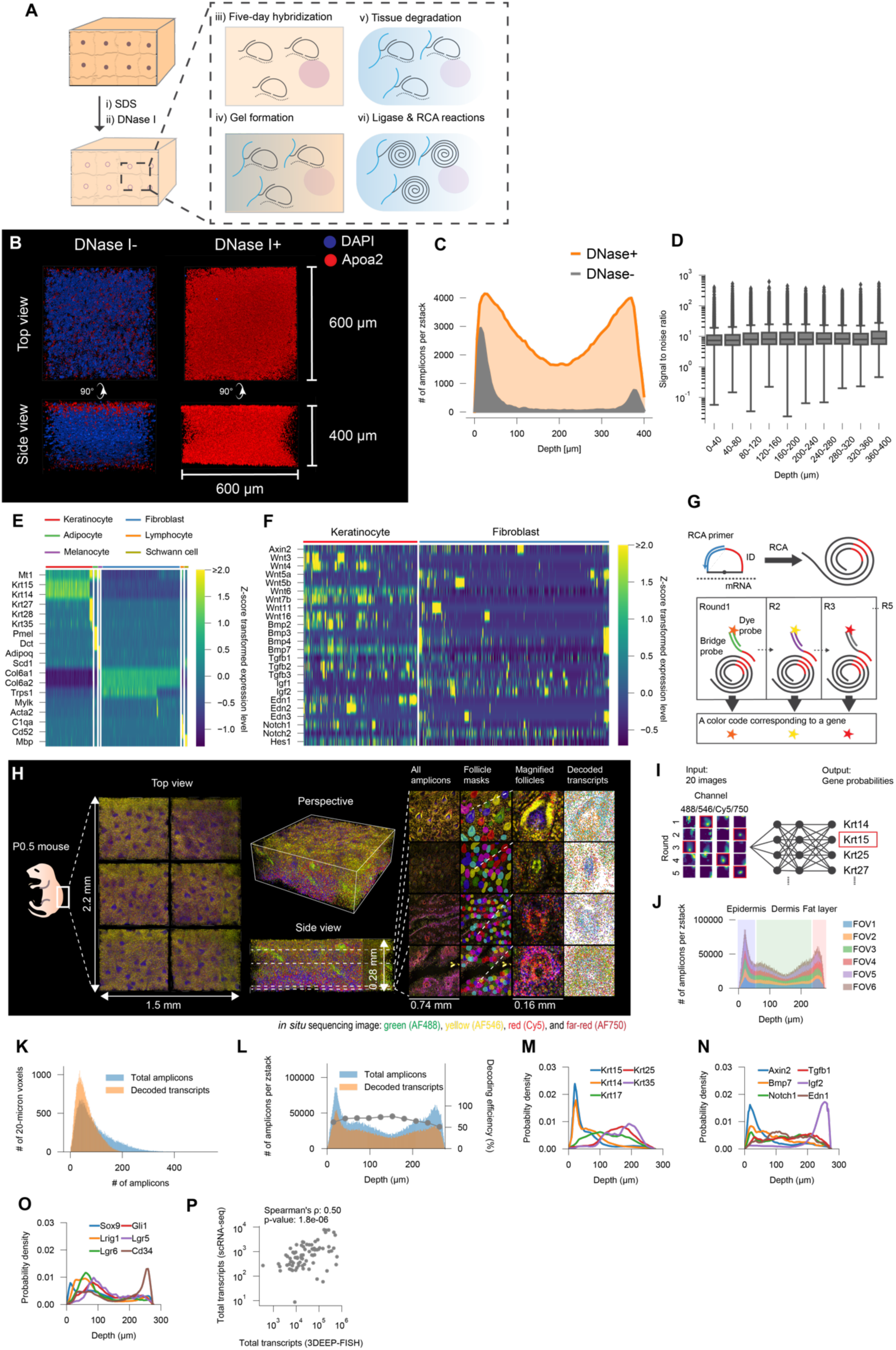
3DEEP combined with FISH for spatial transcriptomics in depths of intact mouse skin. (**A**) Schematic illustration of 3DEEP-FISH approach. Blue lines: hydrogel; dotted lines: RNA transcript, black lines: padlock probe, RCA primer, or RCA amplicon. (**B**) Side-by-side comparison of *Apoa2* mRNA detection in 400 μm-thick blocks of liver with and without DNase I treatment for gDNA removal. gDNA is stained with DAPI (blue) and *Apoa2* amplicons are shown in red. (**C**) Area graphs showing the abundance of *Apoa2* detection across the 400 μm-thick block of liver with and without DNase I treatment. (**D**) Boxplots showing the SNRs of *Apoa2* amplicons across different tissue depths. SNRs were calculated by dividing the signal intensity of amplicons by the standard deviation of the signal intensity within a 7-10 μm radius outer disk surrounding it. (**E**) Heatmap of expression for 18 representative genes in single cells in P0 mouse skin (GSE131498) clustered by cell type, showing enrichment of each gene (rows) in each cluster of cells (columns, labeled based on color key on top) according to the color key on the right. The vertical white bars demarcate cell type clusters. (**F**) Heatmap of expression for 24 morphogen genes in keratinocyte and fibroblasts of P0 mouse skin (GSE131498) clustered by cell type, showing enrichment of each gene (rows) in each cluster of cells (columns) according to the color key on the right. (**G**) Schematic representation of the HybISS decoding process. (**H**) Overview of the 3DEEP-FISH raw data (i.e., amplicons) in intact P0.5 mouse skin, showing six FOVs. Representative images at different depths are shown with colored masks to visualize each hair follicle. The magnified views show the morphological features of hair follicles and decoded transcripts. All transcript images correspond to the second cycle. (**I**) Decoding scheme of 3DEEP-FISH, with images from one representative amplicon in four channels (columns) and five rounds (rows). Detected amplicons were decoded using machine learning based on known correspondence between IDs and genes. (**J**) Stacked bar plots quantifying detected amplicons across skin depth and six FOVs. Each bar corresponds to one of 276 optical sections taken 1 micron apart. (**K**) Histogram showing the total number of detected amplicons and the number of decoded transcripts across 20×20×20 voxels packing the entire scanned volume. (**L**) Barplots showing the number of detected (blue) and decoded (orange) amplicons across skin depth. Black line show decoding efficiency, that is the ratio of decoded amplicons to total amplicons in each scanned optical section, according to the Y-axis on the right. (**M**) Line graphs showing the distribution of select keratin genes across skin depth. (**N**) Line graphs showing the distribution of select morphogen genes across skin depth. (**O**) Line graphs showing the distribution of select stem cell-related genes across skin depth. (**P**) Scatter plot showing the correlation between the abundance of the 81 targeted transcripts between scRNA-seq (GSE131498) and 3DEEP-FISH data across the whole tissue.

### Selecting a gene panel to characterize hair follicle formation

Characterizing hair follicle formation using 3DEEP-FISH requires selecting a panel of marker genes that can identify various cell types in the developing organ. We compiled a set of 85 genes for cell type annotation and functional evaluation of developing hair follicles (**Table 1**). Briefly, we first curated an initial list of marker genes that were independently validated by multiple studies for different hair follicle structures. We then augmented this initial set using a high-quality scRNA-seq dataset from mouse P0 dorsal skin (Ge et al. 2020) to identify other genes showing differential expression between cell types within the skin (**Fig. 1E**, **Table 1**). The chosen genes include 34 for cell type annotation: 13 for keratinocytes subtypes, 8 for fibroblast subtypes, 3 for melanocyte subtypes, and 10 for other cell types residing in the skin (**Fig. 1E**). Figure 1E shows the distinct cellular clusters, namely keratinocyte, melanocyte, adipocyte, fibroblast, lymphocyte, and Schwann cell, separated by the subset of the curated genes. These genes capture the subclusters of keratinocyte and fibroblast clusters. The list also includes 51 genes for functional evaluation: 9 stem cell markers, 40 morphogen-related genes (10 for the *Wnt* pathway, 16 for the *Tgf* superfamily pathway, 5 for the *Notch* pathway, 4 for the *Igf* pathway, 5 for the *Edn* pathway), a chemokine marker, and a marker of bacterial cells. Overall, these 85 genes capture the structural and functional diversity of cells within the hair follicle (**Fig. 1F**).

**Table 1.** List of gene used for characterizing new-born mouse skin. Each gene is listed together with its functional classification, associated cell lineage, associated hair follicle structure, related references, and basis of selection for this study.

### Spatial transcriptome of newborn mouse dorsal skin

To characterize these 85 genes in developing hair follicles, we multiplexed 3DEEP-FISH based on Hybridization-Based In Situ Sequencing (HybISS) (Gyllborg et al. 2020). We designed and synthesized up to five padlock probes for each of the 85 transcripts in our panel (**Table S1**). Each probe contains a 40-nucleotide sequence split into two 20-nucleotide (nt) arms that specifically hybridize to the target transcript, an 18-nt ID sequence that is unique to each transcript and used for downstream identification, and a 19-nt common sequence to prime RCA (**Fig. 1G**). All probes for the same transcript share the same ID. After RCA, the resulting amplicon will contain a concatemer of ID sequences. To decode these IDs with fluorescence imaging, we designed bridge probes and fluorophore-conjugated probes for five sequential rounds of hybridization and imaging in four channels (**Table S2**). Bridge probes are complementary to the amplified ID sequences and one of the four fluorophore-conjugated oligonucleotides. Each round of hybridization uses a different library of bridge probes such that the combination of all rounds labels each ID with a unique permutation of colors over hybridization rounds (**Fig. 1G**). In five rounds of hybridization with four colors, 4^5^ = 1,024 color codes are possible, of which a 144 subset has a Hamming distance of at least two between all pairs. We assigned 85 of these 144 color codes to our transcripts’ IDs, leaving the remaining 59, which do not correspond to any transcript, as negative control ‘ghost IDs’ for estimating decoding accuracy.

Using these probes, we performed 3DEEP-FISH on the dorsal skin of a 12-hour old mouse without slicing the tissue (**Fig. 1H**). We scanned six 0.75 x 0.75 x 0.3 millimeter (X, Y, Z) adjacent fields of view (FOVs) using a spinning disk confocal microscope and a 20X objective, in total covering a 3.3 mm^2^ area of skin at a depth of approximately 300 microns. To analyze raw image stacks and identify amplicons, we initially applied the well-established ExSeq processing pipeline (Alon et al. 2021). The pipeline identified a total of 10,044,124 putative amplicon locations (∼10 rolonies per 10-micron voxel). Of these, 4,771,780 (48%) could be confidently matched to a transcript color code while allowing 5% false positives, as measured by the number of amplicons assigned to ghost IDs. We reasoned that the depth and density of amplicons in our dataset combined with the technical noise unique to each imaging setup prevent the ExSeq pipeline from maximizing data extraction. We thus developed a machine-learning approach based on a convolutional neural network to supplement the pipeline and improve decoding accuracy (**Fig. 1I**). To train the network, we used the images (4 colors by 5 cycles) for the 744,696 amplicons in the top 10% of confidence scores together with their gene calls from the ExSeq pipeline. No ghost IDs were detected among these top 10% amplicons. After training, we input the images for each of the 10,044,124 amplicons into the model to predict gene names. The model returned a likelihood of each amplicon corresponding to each gene, which was used as a confidence score for gene calling. For each gene, we chose a confidence score cutoff across the dataset such that the average ghost ID detection rate was 5% or below. Four transcripts (*Crabp1*, *Ccl2*, bacterial rRNA, and *Wnt7b*) that did not reach this confidence threshold were eliminated from further analysis. Overall, our machine learning approach improved the number of amplicons decoded with high confidence to 6,601,822 (66%), resulting in 6.5 rolonies per 10-micron voxel across approximately 1 mm^3^ of skin.

We analyzed the overall spatial characteristics of the amplicons’ distribution. Amplicon detection efficiency was highest near the two surfaces of the tissue block, as observed in the liver (**Fig. 1C**), but was otherwise consistent across the depth of the tissue and different fields of view (FOVs) (**Fig. 1J**). The density of detected amplicons across the tissue was consistent, with an average of 92 per 20-micron voxel (25th percentile: 43, 75th percentile: 128) (**Fig. 1K**). Decoding efficiency, the fraction of initial amplicons decoded confidently, showed a consistent distribution across both tissue depth and 20-micron voxels (**Fig. 1K,L**). Overall, these results suggest that 3DEEP-FISH performs consistently across different tissues and in a multiplexed setting.

While the total transcript distribution was consistent across space, individual transcripts exhibited spatial heterogeneity. Different transcripts were enriched at varying depths of the tissue (**Fig. 1M–O**). For example, *Krt14* and *Krt15*, which are highly expressed in epidermis, are enriched near the skin surface, while *Krt35*, a hair shaft marker, is enriched deeper in the skin (**Fig. 1M**) (*37*, *38*, *92*). Despite this spatial heterogeneity, the total abundance of each transcript matched expectations: our spatial transcriptomic data correlates with scRNA-seq in overall gene abundances (Spearman’s ρ = 0.50, p-value < 2e-6) (**Fig. 1P**). Overall, these results indicate that our 3DEEP-FISH dataset of 81 genes accurately reflects the heterogeneous tissue structure, portending its ability to capture the spatiotemporal dynamics of hair development.

### Identifying hair follicles within skin tissue

To focus on hair follicles within the captured skin volume, we manually isolated each follicle. Follicles are visually distinguishable in our data as distinct cylindrical shapes that extend inward from the surface and are enriched in keratin expression (**Fig. 2A**). We defined a region of interest (ROI) for each hair follicle by manually identifying the contours of its cross-section in each optical section. **Figure 2A** shows the contours of eight representative follicles of different sizes across five optical sections. This process identified 371 complete follicles in all six FOVs. 359 follicles at the boundaries of the scanned volume or spanning multiple FOVs were excluded from further analysis due to incompleteness or uncertainty about exact 3D organization, respectively. The overall density of follicles was approximately 186 per square millimeter (**Fig. 2B**). The complete follicles were designated HF001 to HF371 based on their position within the imaged tissue (**Table S3**). The ROIs for each follicle are provided in the supplementary materials. These ROIs allowed us to determine the molecular composition of each follicle in 3D. **Figure 2C** shows a 3D rendering of transcripts within the boundaries of the representative follicles from Figure 2A. On average, we detected 5,586 decoded amplicons per hair follicle, with a range from 424 in HF196, one of the smallest follicles, to 33,151 in HF245, the largest (**Fig. 2D**).

**Figure 2.**
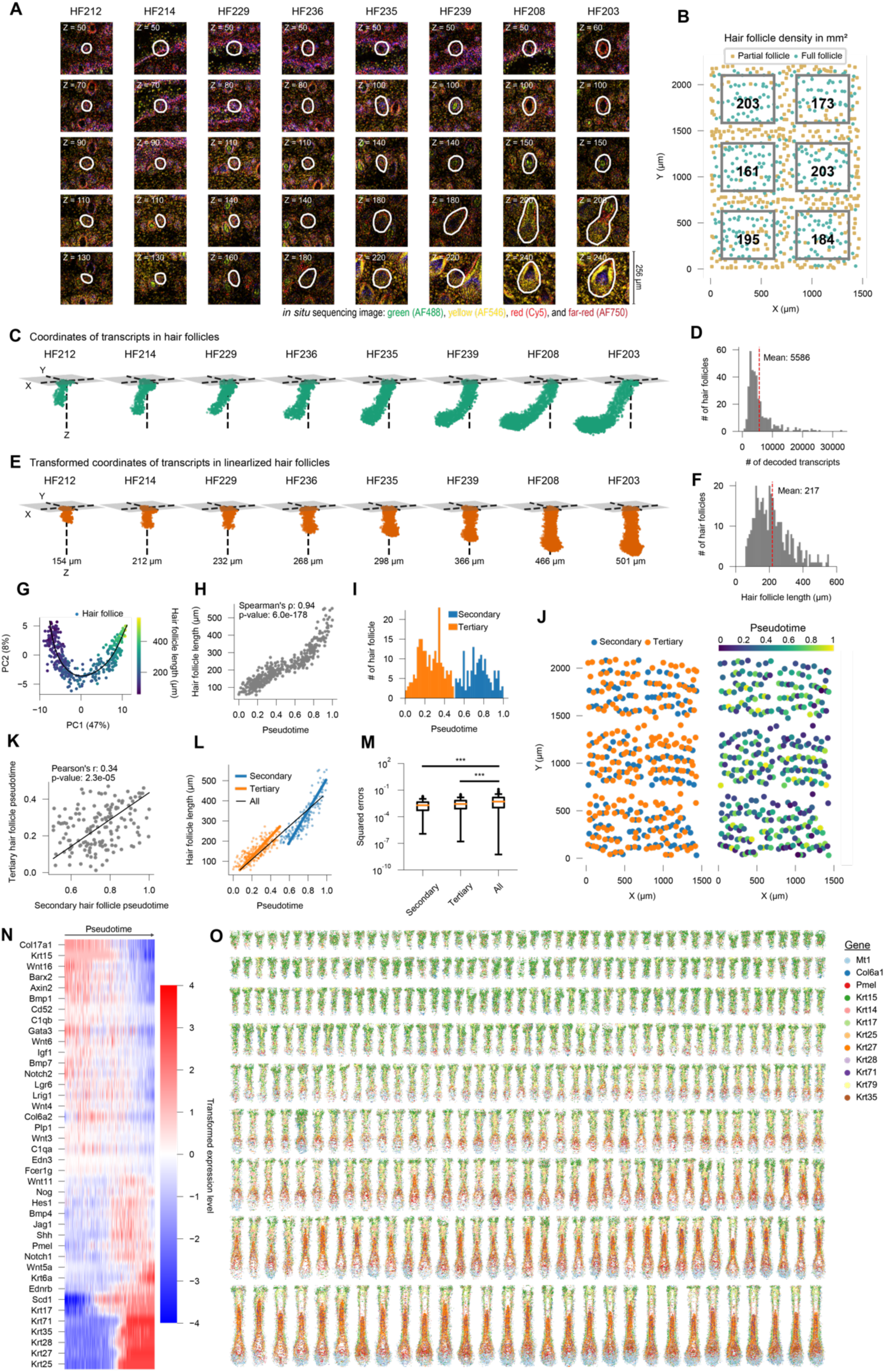
Reconstructing developmental trajectory of hair follicles using whole-organ pseudotime. (**A**) Raw images from eight hair follicles of different sizes at different depths of the skin. Z denotes the Z-stack number of optical image, which approximates depth in microns as optical sections were 1 micron apart. The white line marks the manually generated border for the follicle. (**B**) Scatterplot showing the position of all detected hair follicles across the X and Y axes of the scanned skin area. Full follicles (green) and partial follicles (brown) which cross FOV boundaries are labeled. The rectangles mark the areas within each field of view where hair follicle density was calculated to exclude partial or shared follicle, and the numbers in each represents the density value. (**C**) 3D plots showing transcript coordinates of the follicles in **A**. (**D**) Histogram of the number of decoded transcripts per hair follicle. (**E**) 3D plots showing transcript coordinates for the follicles in **C** after linearization. (**F**) Histogram of hair follicle length after linearization. (**G**) Scatterplot showing the values of Principal Component (PC) 1 and 2 for 371 follicles based on their transcript composition. The color of each point represents follicle length according to the color key on the right. The black line is the Slingshot-fitted trajectory curve used for estimating pseudotimes. (**H**) Scatter plot showing the correlation between pseudotime (from **G**) and length for all hair follicles. (**I**) Histogram of pseudotimes for 371 hair follicles. Color denotes the classification of the follicle as secondary (blue) or tertiary (orange) based on a Gaussian mixture model clustering. (**J**) Scatter plots showing the spatial distribution of hair follicles on the skin surface, colored based on the secondary/tertiary classification (left) or pseudotime (right). (**K**) Scatter plot showing the correlation between the pseudotimes of neighboring secondary and tertiary hair follicle pairs. (**L**) Scatter plot showing the relationship between follicle pseudotime and length with lines fitted for secondary-only (blue), tertiary-only (orange), and all (black) follicles showing improved agreement between pseudotimes and length when tertiary and secondary classes are separated. Fitting was done using the least-squares method. (**M**) Boxplots showing the distribution of squared errors for the linear fits in panel **L**. P-values were calculated using a two-sided Wilcoxon rank sum test. ***: P < 0.001. (**N**) Heatmap showing normalized gene expression for genes (rows) in all hair follicles ordered based on pseudotime (columns). Only the top 40 genes with the most significant correlation are shown. scTransform (Hafemeister and Satija 2019) was used for normalizing expression. (**O**) 2D projection of eight representative genes for all 371 hair follicles, ordered by estimated pseudotime from top left to bottom right. Genes are colored based on the color-code on the right.

Hair follicles do not grow perpendicularly to the skin surface. Moreover, longer follicles in later stages of development may curve or bend. To simplify structural comparisons between hair follicles of different shapes, we developed a computational pipeline to transform each follicle into a linear structure using thin-plate spline transformation while preserving the spatial relationships between transcripts (**Fig. 2E**). This linearization allowed us to measure the overall length of the follicles, which ranged from 57 μm (HF57) to 553 μm (HF245) (**Fig. 2E,F**). Going forward, we will analyze these linearized hair follicles in a cylindrical coordinate system (Z: axial coordinate with 0 at the skin surface and positive values indicating depth within the skin; R: radial distance from the Z-axis; θ: azimuth with 0 along the direction of hair follicle growth), unless otherwise noted. These steps provide a foundation for the molecular analysis of developing hair follicles in 3D.

### Reconstructing developmental trajectory of hair follicles

Based largely on histological characterization, hair follicle formation is classically divided into three broad phases – induction, organogenesis, and cytodifferentiation (Schneider et al. 2009) – which are further broken down into eight stages. During induction (Histological Stages 0 and 1), interactions between adjacent dermal and epidermal cells initiates hair follicle formation. During organogenesis (Histological Stages 2 to 5), the follicle grows, and all its cell types and tissues emerge. During cytodifferentiation (Histological Stages 6 to 8), the follicle matures morphologically, and the hair shaft emerges. Because different follicles develop asynchronously, newborn skin is expected to have follicles spanning all stages of organogenesis and early cytodifferentiation (Paus et al. 1999; Sayama et al. 2010; Saxena et al. 2019). We tested whether molecular profiles of hair follicles reflect their developmental stage. We performed principal component analysis (PCA) on the transcriptome of the 371 follicles by treating each as a homogeneous compartment of transcripts (**Fig. 2G**). The results show that the first two principal components, which explain a 55% of the variation in the dataset, clearly match follicle size, indicating that each follicles’ captured transcriptome reflects its stage along their developmental trajectory. We thus used Slingshot (Street et al. 2018) to place each follicle at a relative position along this developmental trajectory (**Fig. 2G**), thereby creating a whole-organ pseudotime ranging from 0 to 1. Pseudotime 0 here is expected to approximate Histological Stage 2 while pseudotime 1 is expected to approximate Histological Stage 6. Follicle pseudotime startlingly correlates with its length (Spearman’s ρ = 0.94, p-value < 6e-178) (**Fig. 2H**). These results demonstrate that the molecular composition of each hair follicle can be used to infer its developmental stage.

We next analyzed the relationship between follicle developmental stage and spatial arrangement in the skin. Follicle pseudotimes were bimodally distributed, with modes at 0.2 and 0.8 (**Fig. 2I**). This observation is consistent with the well-known waves of mouse hair follicle induction, which occur around E14, E16, and E18, to give rise to primary, secondary, and tertiary follicles, respectively (Duverger and Morasso 2009). Primary, secondary and tertiary hair follicles account for 1–3%, 30% and 65–69% of all follicles, respectively. Therefore, the pseudotime mode at 0.8 largely represents secondary follicles with a minority of primary follicle, whereas the mode at 0.2 likely represents tertiary hair follicles. We thus classified all follicles into simplified groups of secondary and tertiary using Gaussian mixture model clustering of their pseudotimes (**Fig 2I**). When we analyzed the positions of full hair follicles in the skin, we observed regularly spaced rows of follicles in which secondary and tertiary follicles appear to alternate (**Fig. 2J**). Among the full hair follicles, the closest hair follicle to 85% of the secondary hair follicles is a tertiary follicle, which is significantly higher than the 58% that would be expected by chance (Chi-squared test of independence, p-value < 6e-7). Moreover, the developmental stage of all adjacent secondary-tertiary follicle pairs is significantly correlated (Pearson’s R: 0.34, p-value < 3e-5) (**Fig. 2K**). Together, these results support a Turing reaction-diffusion model of follicle induction, which posits that tertiary follicles are induced between secondary follicles as skin growth leads to increased distance between them (Sick et al. 2006; Schlake and Sick 2007).

Separating the secondary and tertiary follicles improved the agreement between hair follicle length and developmental stage, revealing differences between the growth dynamics of secondary and tertiary follicles. The mean squared error of the linear regression between follicle stage and length is 0.011 for all follicles combined but is significantly lower when secondary and tertiary follicles are separated, at 0.005 and 0.004, respectively (Wilcoxon rank-sum test, both p-values < 2e-15) (**Fig. 2L,M**). These results also demonstrate the accuracy of a molecular approach for staging hair follicles compared to the standard approach of measuring length, as the size overlap between the oldest tertiary follicles and the youngest secondary follicles prevents accurately staging them based on size alone.

The molecular staging of the follicles enabled us to characterize the changes in gene expression during hair follicle development (**Fig. 2N**). Hair follicles in the earliest stages of morphogenesis, represented by pseudotimes close to 0, are enriched in subsets of *Wnt* and *Bmp* genes (*Wnt16*, *Wnt6*, *Axin2*, and *Bmp1*), whereas those in the later stages of morphogenesis, represented by pseudotimes close to 1, exhibit elevated *Notch* signaling genes (*Jag1* and *Hes1*) (**Fig. 2N**). Additionally, a transition from a single keratin gene to a diverse set (*Krt15* to *Krt17*, *Krt25*, *Krt27*, *Krt28*, *Krt35*, and *Krt71*) was observed, indicating the successful isolation of hair follicles from the early and late stages. More importantly, this molecular staging enabled us to order the 3D structures of hair follicles along their developmental trajectory, creating a 371-frame stop-motion animation of their development in time and space (**Fig. 2O**).

### Identifying the cell types and structures of hair follicles in space and time

The stop-motion animation clearly conveys a visual sense of hair follicle growth, with newly obtained cells and structures marked by the emergence of different keratin genes. To map this structural evolution at a cellular level, we first inferred cell boundaries within each follicle. 3DEEP-FISH, similar to the majority of other spatial transcriptomic methods (Wang et al. 2018; Wang et al. 2021), does not preserve exact cell boundaries. To infer approximate boundaries, we applied Baysor, an expectation maximization (EM) algorithm that iteratively optimizes inferred cell boundaries to maximize transcriptional similarity within each cell (**Fig. 3A**) (Petukhov et al. 2022). To minimize the possibility that distinct cells are mistakenly combined into a single boundary by Baysor, we used a conservative 5-micron expected cell radius. Across all follicles, we obtained a total of 122,059 inferred cells, ranging from 23 (HF196) to 1,826 (HF245), with the total number of cells correlating with follicle size (**Figs 3B, C**). Henceforth, we will refer to these inferred cells simply as cells. The average cell contains 16 amplicons (**Fig. 3D**); the average transcript is covered 0.19 times per cell, which is threefold higher than the number of UMIs per transcript per cell for the same set of genes in matching scRNA-seq data (**Fig. 3E**) (Han et al. 2018).

**Figure 3.**
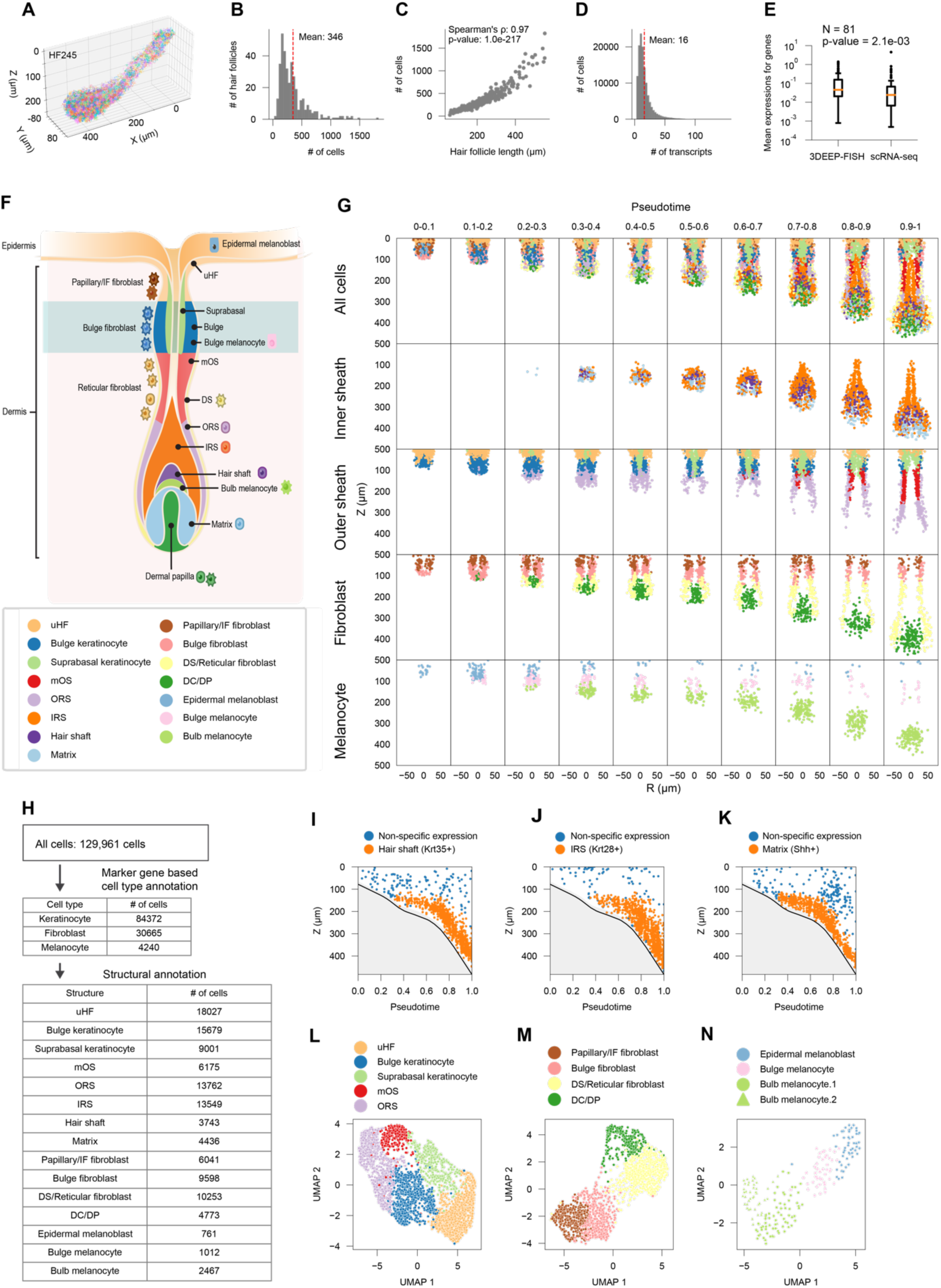
Annotating cell types and structures within hair follicles. (**A**) 3D image of all HF245 transcripts colored by their single-cell assignment based on Baysor. (**B**) Histogram of the number of inferred cells per hair follicle. (**C**) Scatter plot showing the correlation between hair follicle lengths and their number of cells. (**D**) Histogram of the number of transcripts per cell. (**E**) Box plots comparing the number of unique transcripts detected per cell between 3DEEP-FISH and scRNA-seq (GSE108097) for the shared gene set. P-value was calculated using Wilcoxon signed-rank test. (**F**) Schematic illustration of hair follicle, with various cell types and structures identified according to the color key on the bottom (by Jennifer E. Fairman, CMI, FAMI, © 2024 JHU AAM). (**G**) Scatter plot showing the spatial coordinates of hair follicle cell types and structures faceted by cell type groups (rows) and pseudotime bins. Color key the same as panel **F**. For each pseudotome bin, all hair follicles within that bin were overlayed and a cross section is shown. (**H**) Cell type/structure annotation workflow with the number of cells at each step. (**I**) Scatter plot of the spatiotemporal coordinates (pseudotime and axial position Z) of *Krt35*+ keratinocytes. *Krt35* marks the hair shaft. Density-based clustering was used to separate the main hair shaft cluster of cells (orange) from those expressing it sporadically in the background. (**J**) Scatter plot of the spatiotemporal coordinates of *Krt28*+ keratinocytes which mark the IRS. Density-based clustering was used to separate the main IRS cluster (orange) cells expressing it sporadically in the background. (**K**) Scatter plot of the spatiotemporal coordinates of *Shh*+ keratinocytes. *Shh* marks the matrix. Density-based clustering was used to separate the matrix cluster (orange) cells expressing it sporadically in the background. (**L–N**) UMAP scatter plot of spatiotemporal pseudobulks for outer sheath keratinocytes (L), fibroblasts (M), and melanocytes (N). Colors signify the different clusters as identified using Leiden and based on the key on top.

We assigned these cells to different cell types and structures based on their transcript composition and spatiotemporal positions in the follicle (**Fig. 3F–H**). We began by classifying cells into fibroblast, melanocyte, and keratinocyte lineages based on their marker genes, as listed in Table S1 (**Fig. 3G,H**). This procedure placed 98.8% of all cells in one of these three lineages; the remaining 1.2% were excluded from further analysis. We next assigned cells in each lineage to anatomical structures in hair follicles. While the three main lineages have mutually exclusive markers, making their identification straightforward (**Fig. 1E**), their substructures can overlap in markers, requiring simultaneous consideration of marker expression, anatomical position, and hair follicle stage (Joost et al. 2016; Joost et al. 2020). Firstly, we classified the keratinocytes that comprise the inner structures of the follicle because they have the most distinguishable markers and occupy contiguous volumes. These structures include the hair shaft at the center, the inner root sheath (IRS), which surrounds the hair shaft, and the matrix, located at the base of the bulb (**Fig. 3F**). To identify these three structures, we placed all keratinocytes that expressed the structures’ specific marker genes (**Table S1**) in a three-dimensional space with axes corresponding to their axial position (Z) in their follicle, radial position (R) in their follicle, and their follicle’s pseudotime (Methods). We then used density-based clustering in this 3D space to classify keratinocytes belonging to each of the hair shaft, IRS, and matrix (**Fig. 3I–K**). Secondly, we classified the remaining keratinocytes, which constitute the outer sheath that envelops the follicle from the skin down to the bulb. These cells tend to be similar in transcriptional profiles but show slightly different transcriptomes based on their positions within the follicle (Joost et al. 2016; Joost et al. 2020). We first made pseudobulks of these outer sheath keratinocytes by fusing spatiotemporally proximal cells. This process generated 3,160 representative pseudobulks with on average 20 cells and 354 amplicons each (Methods). We applied network-based Leiden clustering (Traag et al. 2019) to the pseudobulks’ transcriptional profiles to identify five clusters (**Fig. 3L**). Based on the agreement between their transcriptional profiles and previous work classifying the outer sheath keratinocytes (Sennett et al. 2015; Joost et al. 2016; Joost et al. 2020), we classified keratinocytes in each cluster as outer root sheath (ORS), mid-part outer sheath (mOS), bulge, suprabasal, and upper hair follicle (uHF) (**Fig. 3G,L**). These five regions of the outer sheath occupy different depths in the follicle, from the ORS, which wraps the bulb, to the uHF, which connects to the skin surface (**Fig. 3F,G**). Thirdly, we classified all the fibroblasts in and around the hair follicle. We employed a similar spatiotemporal pseudobulking approach as we did for the outer sheath keratinocytes, generating 1,557 representative pseudobulks with on average 20 cells and 251 amplicons each. We applied network-based clustering to these pseudobulks, which identified four classes of fibroblasts in and around the hair follicle (**Fig. 3M**): (1) the dermal condensation or dermal papilla (DC/DP) fibroblasts; dermal condensation induces hair formation at early stages and, over time, migrates inside the bulb to form the dermal papilla; (2) dermal sheath or reticular (DS/Reticular) fibroblasts, which wrap the outer sheath keratinocytes in the reticular layer of the dermis; (3) bulge fibroblasts, which are situated above the DS/Reticular fibroblasts and wrap the bulge keratinocytes; and (4) papillary and inter-follicular (Papillary/IF) fibroblasts, which wrap the outer sheath keratinocytes in the papillary layer of the dermis and form the top fibroblast layer beneath the skin surface (**Fig. 3F,G**). Lastly, we classified all melanocytes. We again employed the spatiotemporal pseudobulking approach, generating 212 representative pseudobulks with on average 20 cells and 327 amplicons each. We applied network-based clustering to these pseudobulks which identified four groups of melanocytes in the hair follicle (**Fig. 3N**). Based on spatial location, we classified the cluster on skin surface as “epidermal melanoblasts”, the cluster at the hair bulge as “bulge melanocytes”, and the two clusters in the hair bulb as “bulb melanocytes” (**Fig. 3F,G**). The bulge melanocytes can be considered the same as melanocyte stem cells in later stages of development, whereas the bulb melanocytes can be considered mature melanocytes (Zhang et al. 2023).

### Cellular dynamics of hair follicle formation

The classification of hair follicle cells into its various structures enabled measuring the growth rate of each structure and cell type over time (**Fig. 4A–O**). The results show the cascades of cell type emergence from all three lineages during hair follicle organogenesis (**Fig. 4P**) with three distinct stages. During the first stage, between pseudotimes 0 and 0.2, the follicle is dominated by uHF, bulge keratinocytes, papillary/IF fibroblasts, bulge fibroblasts, epidermal melanoblasts, and bulge melanocytes. In the second stage, between pseudotimes 0.2 and 0.7, cell type complexity of the follicle increases with the formation of the bulb and the ordered emergence of suprabasal keratinocytes, ORS, bulb melanocytes, matrix, hair shaft, and IRS, followed by late induction of mOS. In the final stage, between pseudotimes 0.7 and 1, the follicle transitions into a growth phase, with rapid expansion of mOS, hair shaft, matrix, ORS, IRS, DS/Reticular fibroblasts, and DC/DP fibroblasts. The behavior of different cell types is distinct during these windows. The uHF, papillary/IF fibroblasts, and epidermal melanoblasts show steady numbers throughout. The bulge keratinocytes, bulge fibroblasts, and bulge melanocytes show a burst early in follicle development (0-0.2) and remain steady in number thereafter. DS/Reticular fibroblasts, DC/DP fibroblasts, suprabasal keratinocytes, matrix, hair shaft, ORS, and bulb melanocytes initially increase between 0.2-0.4 and show a second burst in number after 0.7. IRS emerges around 0.3 and continues to grow throughout to become the most abundant cell type in the hair follicle by pseudotime 1. Finally, mOS emerges around 0.6 and expands rapidly after 0.8. These observations capture the complicated dynamics of cell type emergence followed by growth during hair organogenesis. They also update the histological descriptions of hair follicle formation (Paus et al. 1999; Saxena et al. 2019; Welle 2023). Whereas these analyses had suggested that matrix and IRS emerge first, followed by ORS, and then hair shaft, our results clearly show that these populations emerge almost concurrently in a short window of time during organogenesis at pseudotime 0.2–0.3, with the ORS appearing first (**Fig. 4P**). A possible explanation for this discrepancy is that matrix and hair shaft cells remain small in numbers after initial emergence and only expand in the later stages of development. Therefore, histological analyses may have only been able to identify these cells after they grow in numbers. Another possibility is that molecular changes accompanying the differentiation of these cells precede the histological changes, making it challenging for histological approaches to identify their presence in the earliest stages of their differentiation.

**Figure 4.**
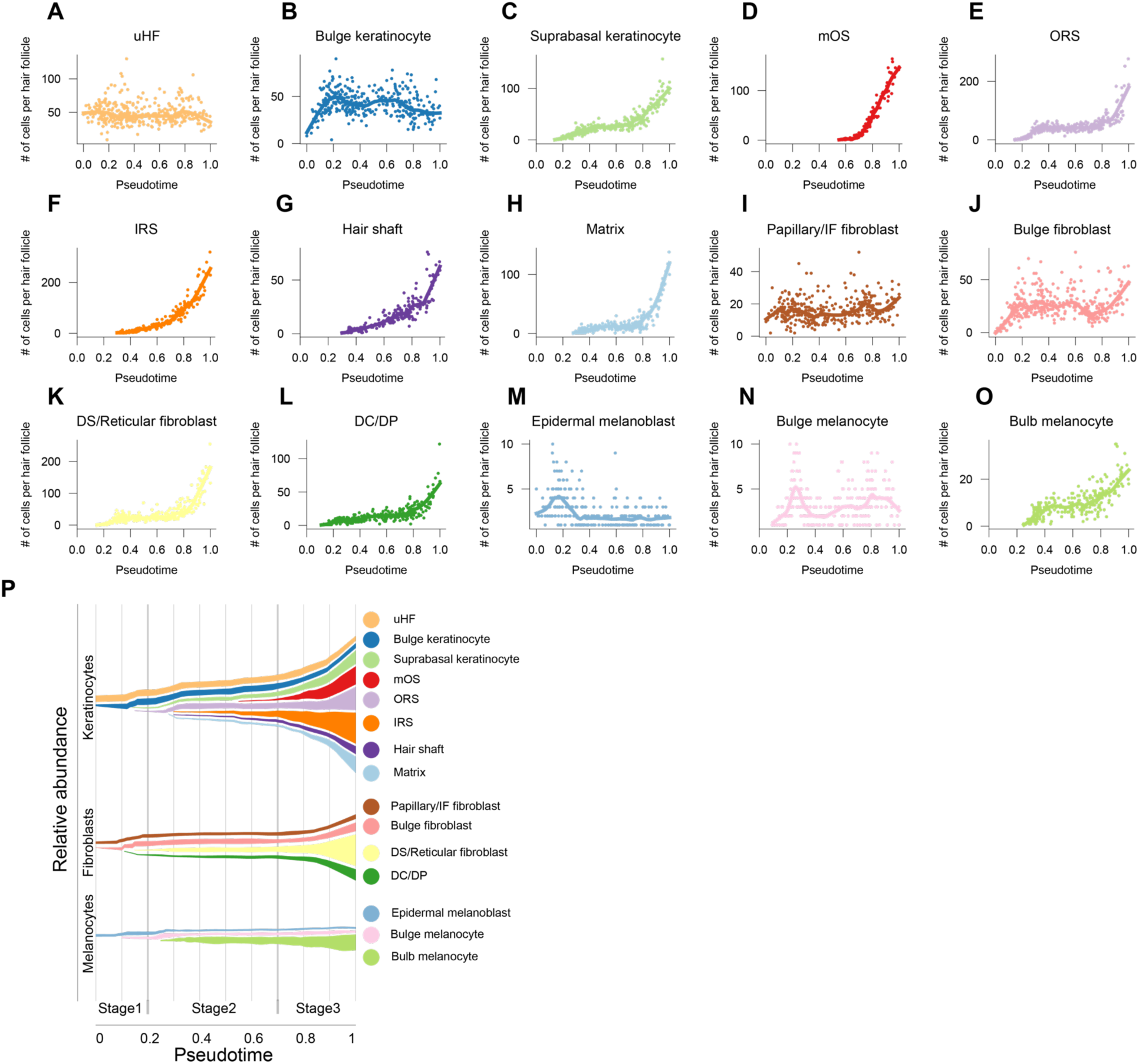
Cell population dynamics during hair follicle development. (**A-Q**) Scatter plots of each cell type’s count per follicle over pseudotime. Corresponding cell type is labeled on the top each panel. Colors are based on Fig. 3F. Locally weighted scatterplot smoothing (LOWESS) is shown as solid line. (**P**) Stream plot summarizing the growth of different cell populations in hair follicles over pseudotime.

### Spatiotemporal dynamics of morphogens

Our gene panel of 81 genes included 40 morphogens and morphogen receptors. These genes represent the changes in the extrinsic environment that the cells in the hair follicle establish and experience. We created a spatiotemporal atlas of changes in each morphogen and receptor in the hair follicle (**Fig. S1**) and all annotated structures (**Figs. S2 and S3**). These data show the complex dynamics, in space and in time, of morphogen expression throughout hair follicle organogenesis involving all five targeted morphogen families, namely the *Wnt* family, the *Tgf b* superfamily, the *Notch* family, the *Igf* family, and the *Edn* family. However, a few distinct patterns emerge. First, fibroblasts and keratinocytes show lineage-specific morphogen profiles even when in spatial proximity. We divided the pseudotime window into nine bins and averaged the expression of all cells from the same hair follicle structure within each bin to calculate overall expression level (z-score) and significance (q-value) for each morphogen over time (**Fig. 5A**). The results show a more similar morphogen profile between the same cell lineage in different structures than between different cell lineages (**Fig. 5A**). For example, in both the upper and lower halves of the hair follicle, *Nog* and *Igf1* are enriched in fibroblasts, whereas *Hes1* is enriched in keratinocytes (**Fig. 5A**). This observation indicates that the distinct functional and structural roles of different lineages are also reflected in their signaling profiles. Second, morphogen patterns are more stable over time in the upper region of the hair follicle compared to the lower region of the hair follicle. For example, uHF keratinocytes and bulge fibroblasts maintain their overall morphogen profiles over time (**Fig. 5A,B**). By contrast, DC/DP and matrix show extensive temporal dynamism in their morphogen profiles (**Fig. 5A,B**). We quantified this dynamism by calculating the variance of z-scores over pseudotime for the enriched morphogen genes in each structure (**Fig. 5C**). The result indicates that uHF morphogen expression is the most stable over time with the lowest variance in mean expression levels, while lower part structures, including DC/DP, DS/Reticular fibroblasts, and ORS, show the highest variations in morphogen gene expression. When average variance corresponding to each structure was grouped for upper and lower structures, the overall variance was significantly higher in the lower part (**Fig. 5D**). This contrast in morphogen dynamism between the upper and lower halves of the follicle is reflected in their structures as well. For example, the papillary/IF and bulge fibroblasts in the upper parts of the follicle occupy stable locations that minimally change over time (**Fig. 5E**), whereas DC/DP, IRS, and matrix in the bulb region show extensive structural changes over time (**Fig. 5E**). This contrasting behavior of the upper and lower hair follicle during development matches the mature hair follicle wherein the upper part does not change visibly and the lower part is continuously remodeled during the hair life cycle (Schneider et al. 2009). Therefore, these results suggest that the foundation for the divergent behavior between the upper and lower parts of the hair follicle in the adult stage is laid during ontogeny. Third, a signaling hub is distinguishable on each end of the follicle. In the upper part of the hair follicle, uHF keratinocytes are enriched in 12 out of the 40 morphogen-related genes tested across the developmental window; no other upper-half structure is enriched in more than 7 (**Fig. 5B**). In the bulb region, DC/DP fibroblasts are enriched in 22 different morphogen-related genes at some point in the developmental window, which is the highest among all structures (**Fig. 5B**). DC/DP is well-known as a dynamic signaling niche during hair follicle formation and cycling. Our results capture DC/DP’s dynamism as a shift from *Bmp3*, *Bmp4*, *Bmp7*, *Dll1*, and *Igf1* signaling genes to *Gdf10*, *Igf2*, *Nog*, *Tgfb2*, and *Wnt5a* over the course of hair follicle organogenesis. Taken together, these results are consistent with a model in which the stable upper hair follicle is maintained by a stable signaling center in uHF keratinocytes, and the dynamic lower follicle is guided by a dynamic signaling center in DC/DP.

**Figure 5.**
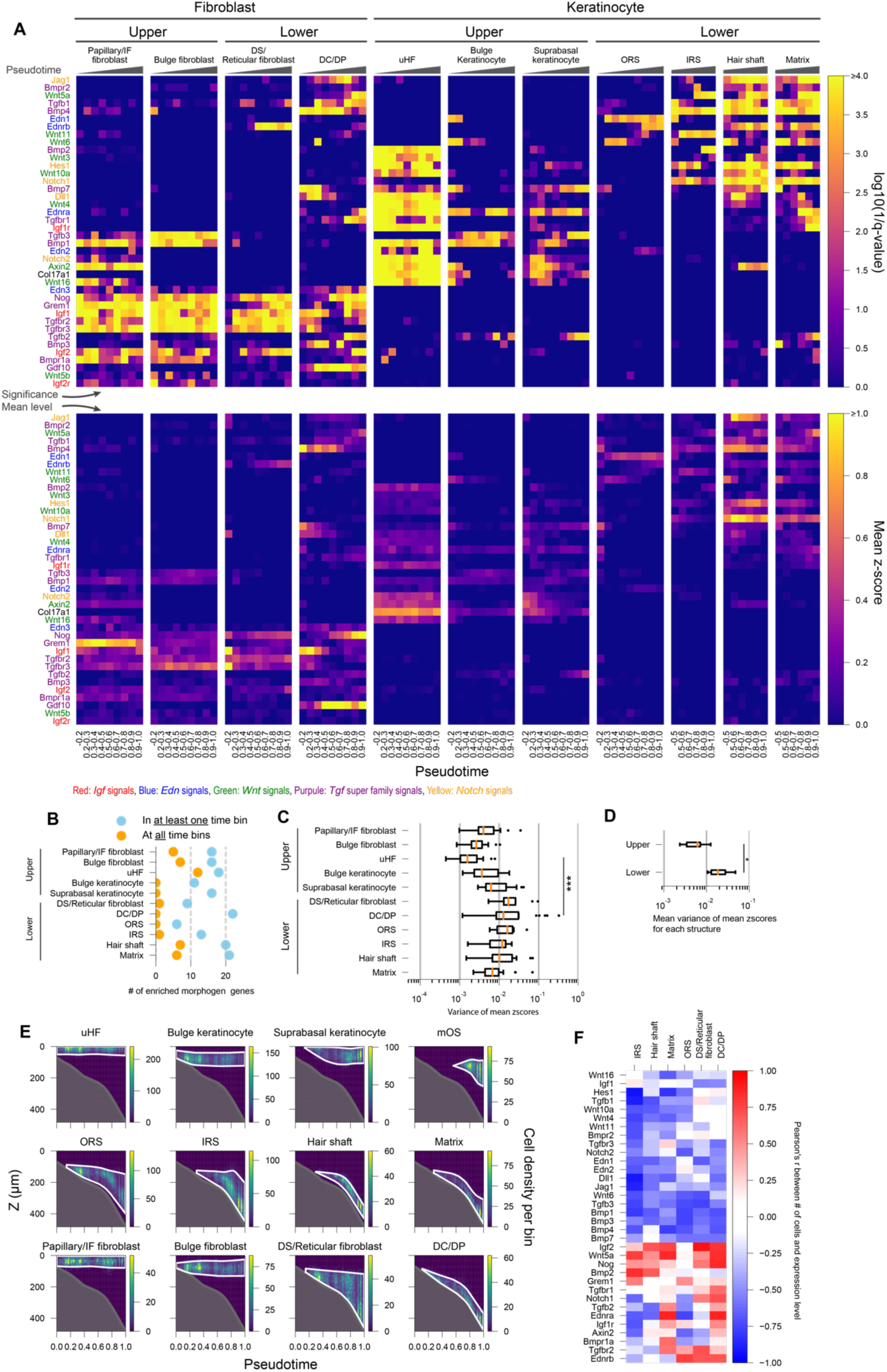
Morphogen expression over space and time. (**A**) Top: Heatmaps of q-values for enriched morphogen genes (rows) across pseudotime bins (columns) faceted by different hair follicle structure. Log10(1/q-value) was capped at 4 to facilitate visualization. Bottom: Heatmaps of z-scores of log-transformed mean expression levels for enriched morphogen genes (rows) across pseudotime bins (columns) faceted by hair follicle structure. Z-score values were capped at 1 to facilitate visualization. Gene labels are colored in red for *Igf* pathway, green for *Wnt* pathway, blue for *Edn* pathway, purple for *Tgf* superfamily pathway, and yellow for *Notch* pathway. (**B**) Cleveland Dot Plots of the number of significantly enriched morphogens genes for different hair follicle structures. Blue: number of genes enriched in at least one pseudotime bin in panel **A** with q-value < 0.05; Orange: number of genes enriched in all pseudotime bins in panel **A** with q-value < 0.05. (**C**) Boxplots showing the variance of mean Z-scores across pseudotimes from panel **A** for enriched morphogens in each structure. Structures are ordered based on being in the upper or lower parts of the follicle. P-values from two-sided Wilcoxon rank sum test. ***: P < 0.001. (**D**) Boxplot showing the average variance of mean Z-scores for all upper and lower hair follicle structures calculated from the values in panel **D**. P-values from two-sided Wilcoxon rank sum test. * : P < 0.05. (**E**) Heatmaps showing the density of each cell type/structure in hair follicles over pseudotime and along the hair follicle’s axial coordinate Z faceted by the cell type/structure. The pseudotime and Z position were split into 50 bins each. Color key for cell density per bin is shown to the right of each plot. (**F**) Heatmap of Pearson correlation coefficients between the number of cells and mean expression level over pseudotime for each morphogen gene (rows) in seven different hair follicle structures (columns).

The changes in morphogen expressions were not restricted to DC/DP. Hair follicle structures that grow significantly over time show more extensive changes in morphogen expression. We identified these changes by calculating the correlation between the number of cells in each structure and the expression levels of each morphogen within that structure (**Fig. 5F**, **Figs. S1– 3**). Many growing structures, namely IRS, hair shaft, matrix, ORS, DS/Reticular fibroblasts, and DC/DP, show either a negative or a positive correlation between cell numbers and morphogen expression for most morphogens (**Fig. 5F**), capturing a shift from a set of signaling pathways (i.e., the negatively correlated) to another set (i.e., the positively correlated) over time and upon growth. For example, the formation and growth of IRS involves a shift from *Notch* signals (see *Notch2*, *Hes1*, *Jag1*, *Dll1* in **Fig. 5C**), canonical *Wnt* signaling (*Axin2*), and a subset of *Bmp* signaling (see *Bmp1*, *Bmp3*, and *Bmp4*) to *Bmp2*, *Wnt5*, and *Noggin* (**Fig. 5F**). These results are unique in mapping a large number of morphogen genes over space and time during the development of an organ. They capture the broad and dynamic changes in morphogen signaling and extrinsic cues during organ development that accompany differentiation, growth, and morphogenesis.

### Formation of the hair follicle canal and opening

The formation of the hair canal inside the follicle is a drastic morphological transformation during its development that prepares it for its eventual function – hair production. Most histological annotations place the hair follicle opening in Stage 6 (Paus et al. 1999; Saxena et al. 2019; Welle 2023), after the end of hair organogenesis. However, the structural changes associated with this process are poorly understood. Mutant mice incapable of forming the hair shaft still form the canal and opening of the follicle, demonstrating that these events are independent of hair shaft growth (Vidal et al. 2005; Nowak et al. 2008). Instead, canal and opening formation are believed to involve rearrangement and apoptosis of *Krt79*-positive (*Krt79*+) keratinocytes at the center of the upper hair follicle (HARDY 1949; PINKUS 1958; Mesler et al. 2021). We analyzed raw 3DEEP-FISH images around the epidermis and observed openings, or ostia, in late-stage follicles (**Fig. 6A**). We then measured the fraction of open follicles over time. The results show that follicles open in the 0.65–0.85 pseudotime range: no follicles were open before 0.65, half were open between 0.7–0.8, and all were open after 0.85 (**Fig. 6B**). We then evaluated the dynamics of this opening process by measuring the radial distribution of *Krt79*+ cells at the skin surface in pseudotime 0.5–1.0. The results show that the diameter of the opening grows at a steady rate between 0.7 and 0.9 (**Fig. 6C**); after 0.9, follicles reach a maximal opening diameter of ∼25 microns.

**Figure 6.**
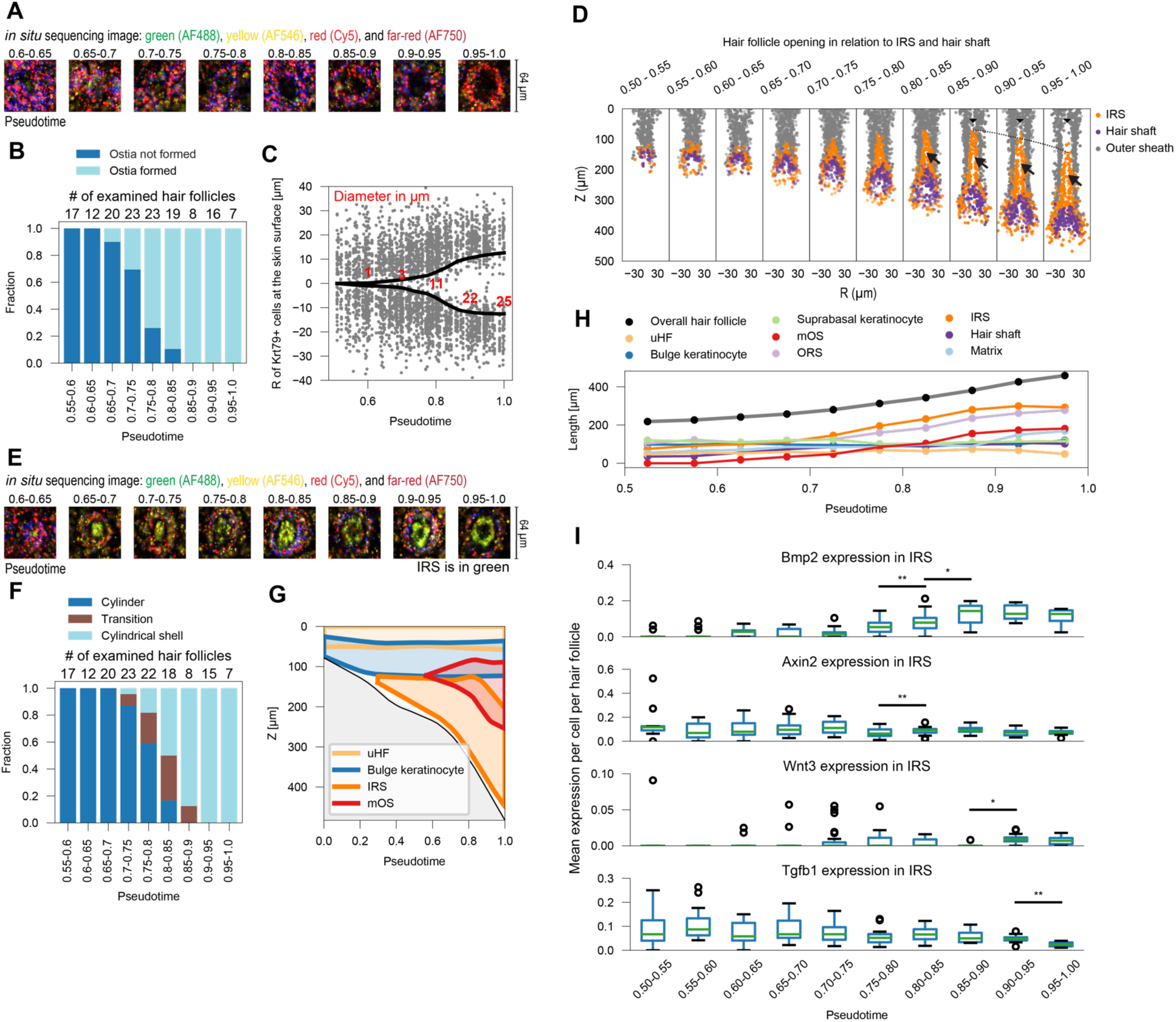
Structural changes leading to hair follicle ostia and canal formation. (**A**) Raw fluorescence images from eight hair follicles, each representing the pseudotime range denoted above the image, near the skin surface, depicting ostia and canal formation. The four colors correspond to fluorescence channels (AF488, AF546, Cy5, and AF750) and correspond to the second round of imaging. (**B**) Stacked barplots showing the fraction of open (light blue) and closed (dark blue) hair follicles in each pseudotime bin. (**C**) Scatter plot showing the distribution of *Krt79+* cells at the hair follicle surface along the radial axis (R) and over pseudotime. *Krt79+* cells from all follicles within the pseudotime window are overlayed. Black lines mark the receding edge of the opening and red numbers quantify its diameter in microns. (**D**) Scatter plots of the axial (Z) and radial (R) positions of IRS (orange), hair shaft (purple), and outer sheath keratinocyte (grey) cells in hair follicles, faceted by pseudotime range. For each facet, all follicles within the pseudotime range have been overlayed. Wedges show the opening of *Krt79+* keratinocyts; arrows mark the opening of the IRS; horizontal dashed lines mark the top edge of the IRS and highlight its downward movement. (**E**) Raw fluorescent images from eight hair follicles, each representing the pseudotime range denoted above the image, showing transformation of the IRS into a cylindrical shell. The four colors correspond to fluorescence channels (AF488, AF546, Cy5, and AF750). The image is from the second round of hybridization where IRS markers (*Krt27* and *Krt71*) are labeled in green (AF488). (**F**) Stacked barplots showing the fraction of hair follicles with their IRS in a cylindrical (dark blue) or cylindrical shell (light blue) configuration or in transition between the two states (brown) in each pseudotime bin. (**G**) Region plot showing the positions and boundaries of uHF (light orange), bulge keratinocytes (blue), IRS (orange), and mOS (red) in axial position and pseudotime. (**H**) Line charts of length over pseudotime for various hair follicle structures as denoted in the color key on top. (**I**) Box plots of morphogen gene expression level changes associated with ostia and IRS cylindrical shell formation over pseudotime. P-values from two-sided Wilcoxon rank sum test. *: P < 0.05, **: P < 0.01.

To visualize the morphological changes associated with opening and canal formation, we divided the pseudotime range between 0.5 and 1.0 into ten bins of 0.05 and overlaid hair follicle structures within each bin, focusing on keratinocytes around the emerging canal (**Fig. 6D**). The results show that the hair canal and opening form before the hair shaft grows into the canal. This opening process involves three structural rearrangements in rapid succession. First, the center of the uHF starts opening in pseudotime 0.7 (**Fig. 6A,C**). Second, IRS gradually transforms from a cylinder into a cylindrical shell, with the transition occurring most rapidly at pseudotime 0.8 and the inner diameter of the shell reaching approximately 15 microns by pseudotime 1 (**Fig. 6D–F**). Third, IRS starts to move downwards into the dermis after pseudotime 0.8 (**Fig. 6D,G**). This downward movement results in opening of the hair follicle below the bulge area (**Fig. 6D**). It is not due to the IRS shrinking – IRS continues to grow in length during this time (**Figs 4F and 6H**); rather, the growth of the outer sheath, in particular its ORS and mOS regions, results in the overall hair follicle lengthening more rapidly than IRS, thereby pushing it downward into the dermis (**Fig. 6G,H**). Overall, these results suggest that hair follicle ostia and canal formation start from its top, at the epidermal level, and propagate downward towards the emerging hair shaft.

We also analyzed the molecular changes associated with hair follicle opening and canal formation. IRS shows a significant increase in *Bmp2* and decrease in *Axin2* expression around pseudotime 0.75 (Wilcoxon rank-sum test, p-value < 0.01) (**Fig. 6I**). Soon after, IRS undergoes a significant reduction in the expression of the *Wnt3a* (p-value < 0.05) (**Fig. 6I**). These changes appear to coincide with the IRS transformation from a cylinder into a cylindrical shell. Finally, near the conclusion of its structural transformation at pseudotime 0.9, IRS undergoes a significant reduction in *Tgfb1* expression (p-value < 0.05)(**Fig. 6I**). Overall, these results identify signaling shifts in IRS keratinocytes that coincide with hair follicle canal and ostia formation.

### Tracing melanocytes stem cells, keratinocyte stem cells, and their niche

Mature hair follicles regenerate throughout the lifespan, undergoing cycles of regression (catagen), quiescence (telogen), and regrowth (anagen). Their ability to regrow requires resident stem cells, which are quiescent during most of their life cycle but are activated at the outset of anagen to create progenitors of the growing follicle. Two main stem cell populations have been characterized in the hair follicle (Lee and Choi 2024). First, melanocyte stem cells, which regenerate the mature melanocytes that pigment the hair shaft. Second, keratinocyte stem cells, which regenerate various keratinocyte-derived structures (**Fig. 3F**). Both stem cell populations reside in the bulge region of the follicle, which does not undergo destruction during catagen (Ito et al. 2004). We sought to leverage our spatiotemporal map of hair organogenesis to investigate the origins of these two stem cell groups and their niche.

Cells of melanocyte lineage reside in three locations within the follicle at the latest stage of organogenesis captured in our data: in the epidermis, in the bulge, and in the bulb (**Figs. 3F,G and 7A,B**). Melanocytes that reside in the bulge are the stem cells (Nishimura et al. 2002). Tracking the history of these cells back in time to the earliest stages observed in our data provided insights into their history. Before pseudotime 0.2, cells of melanocyte lineage are uniformly distributed throughout the hair follicle (**Fig. 7A,B**). This pattern is consistent with their migration from the epidermis into the follicle (Vandamme and Berx 2019). However, after pseudotime 0.2, these cells stabilize into their eventual locations in the bulge and the bulb, signaling an end to active migration and separation of their fate from migratory melanoblasts into quiescent melanocyte stem cells in the bulge and proliferative mature melanocytes in the bulb. Consistent with this conclusion, we observe proliferation of bulb melanocytes after pseudotime 0.4, whereas bulge melanocytes remain steady in numbers (**Fig. 4N,O**). Overall, these results indicate that future melanocyte stem cells reach the bulge area prior to pseudotime 0.2, which precedes the genesis of lower follicle structures (i.e., IRS, matrix, hair shaft). During the formation of bulb structures, these melanocyte populations separate into distinct bulge and bulb regions.

**Figure 7.**
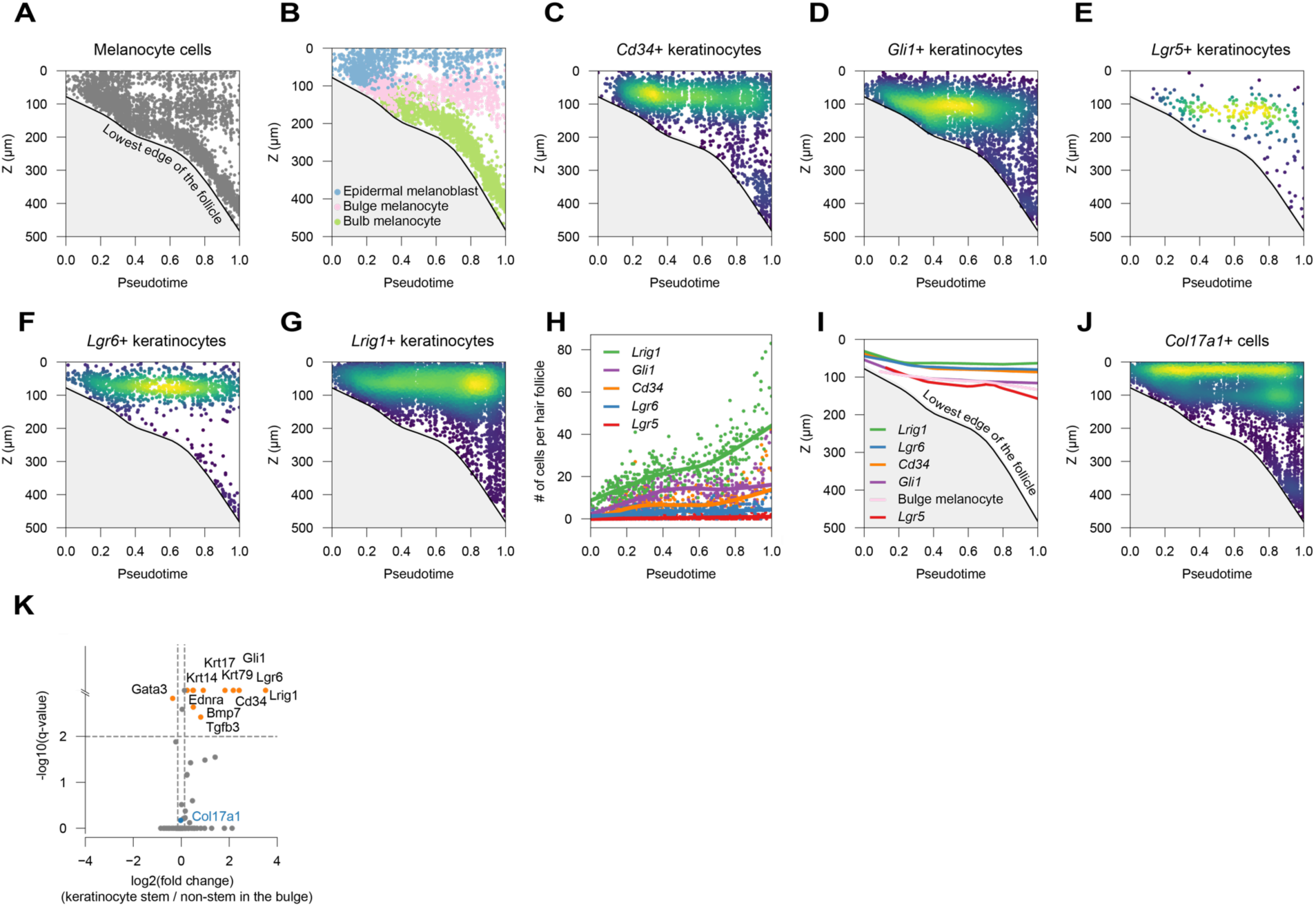
Tracking stem and niche cells during hair organogenesis. (**A**) Scatter plot showing the axial positions of melanocyte cells over pseudotime. Black line marks the lowest edge of follicles at each pseudotime. (**B**) Scatter plot showing the axial positions of epidermal melanoblasts (blue), bulge melanocytes (pink), and bulb melanocytes (green) over pseudotime. (**C-G**) Scatter plots of the axial positions of *Cd34*+ (C), *Gli1*+ (D), *Lgr5*+ (E), *Lgr6*+ (F), *Lrig1*+ (G) keratinocyte stem cells over pseudotime overlayed with their kernel density estimation to visualize the spatiotemporal concentrations of marker gene-positive cells. (**H**) Scatter plot of the number of cells positive for each stem cell marker over pseudotime. Colored lines are LOWESS for each marker. (**I**) Line graphs of the highest-density axial position of keratinocytes positive for different stem cell markers over pseudotime. Local regression was used to estimate stem cell positions in hair follicles. (**J**) Scatter plots of the axial positions of *Col17a1*+ cells over pseudotime overlayed with their kernel density estimation. (**K**) Volcano plot showing up- and down-regulated genes in stem cell marker-positive (*Cd34*, *Gli1*, *Lgr5*, *Lgr6*, and *Lrig1*) keratinocytes in the bulge compared to marker-negative keratinocytes in the same region, with q-values calculated using a two-sided Wilcoxon rank sum test followed by the Benjamini-Hochberg procedure. Log10(1/q-value) values were capped at 3 to facilitate visualization.

Keratinocyte stem cells, similar to their melanocyte counterparts, reside in the bulge area. However, unlike the melanocyte stem cells, they are a heterogeneous population in which different subpopulations are biased toward regenerating different areas of the hair follicle during anagen (Trempus et al. 2007; Jaks et al. 2008; Lee et al. 2008; Nowak et al. 2008; Jensen et al. 2009; Snippert et al. 2010; Brownell et al. 2011; Polkoff et al. 2022). Several marker genes specific to these cells have been described: *Cd34*, *Gli1*, *Lgr5*, *Lgr6*, and *Lrig1*. The inclusion of all these markers among the panel of 81 genes in this study enabled us to characterize their spatial distribution and history (**Fig. 7C–G**). We observed that *Lgr5*+ and *Lgr6*+ stem cells are the least proliferative, suggesting they are quiescent in nature, while *Lrig1+* proliferate rapidly during the capture window of time, suggesting that they may be generating the cell types that nascently emerge during follicle growth (**Fig. 7H**). Interestingly, while all these markers show localization to the bulge area, their exact positions are slightly different (**Fig. 7C–G**). *Lrig1*, *Lgr6*, and *Cd34* localize to the upper bulge area closer to the epidermis, whereas *Gli1* and *Lgr5* localize to the lower bulge area (**Fig. 7I**). While our dataset does not have the resolution to distinguish all stem cell subpopulations, this observation suggests that different keratinocyte stem cell populations are spatially stratified in the bulge. Moreover, all these markers are present in the follicle prior to pseudotime 0.2 (**Fig. 7C–G**), and their relative positions are maintained throughout organogenesis (**Fig. 7I**). These observations suggest that keratinocyte stem cell pools occupy their niche very early in hair follicle formation. We also assessed the spatiotemporal distribution of *Col17a1* (**Fig. 7J**), a stem cell niche marker required for the proper function of both keratinocyte and melanocyte stem cells (Tanimura et al. 2011). *Col17a1*-positive keratinocytes were previously reported in the epidermis and the bulge (Tanimura et al. 2011). Our results recapitulate these reports and further indicate that *Col17a1* starts to be expressed in the bulge before pseudotime 0.2. Taken together, these results suggest that both melanocyte and keratinocyte stem cells, as well as the niche supporting them, are established at the outset of hair organogenesis and maintained thereafter.

We asked whether other genes among our targeted panel are associated with stem cells versus their niche in the bulge. Unlike dissociated scRNA-seq, our data can separate bulge keratinocytes that are positive for the five stem cell markers from those that are negative. The results show that keratinocytes in the bulge that are positive for stem cell markers are most enriched for *Krt79*, followed by *Tgfb3*, *Krt17*, *Bmp7*, *Krt14*, and *Ednra* (**Fig. 7K**). The other keratinocytes in the bulge, which presumably help form the niche for these stem cells, are most enriched for *Gata3*. Enrichment of *Col17a1* is observed in both stem and niche keratinocytes (fold change = 1.0 in stem vs. non-stem). These results suggest that differences in signaling pathway activity exist between stem and non-stem keratinocytes in the bulge region.

## Discussion

In this study, we mapped the development of the most abundant mini organ, the hair follicle, over time and space using a novel spatial transcriptomics technology. We developed a targeted spatial transcriptomic approach, called 3DEEP-FISH, for characterizing tissue blocks as thick as 400 microns. We used this approach to analyze intact newborn mouse skin in which hair follicles develop asynchronously. Through analyzing RNA content of the tissue alone, we captured hundreds of follicles in different stages of formation to create an animated atlas of their development across time and space. Based on this atlas, we characterized the emergence and growth dynamics of different tissues within the hair follicle, mapped morphogen changes over time and space, and dissected structural changes leading to hair follicle opening.

3DEEP-FISH strategy creates a medium that is clear for both imaging and reagent diffusion at depths by casting RNA in a hydrogel using padlock probes and removing genomic DNA along with proteins and lipids (**Fig. 1**). We applied this approach to blocks as thick as 400 microns; however, we expect the approach to work for thicker blocks with increases in diffusion times and reagent concentrations. While we detected target transcripts by implementing a padlock-based strategy (Chen et al. 2018; Gyllborg et al. 2020), the casting and clearing approach of 3DEEP is broadly applicable to *in situ* transcriptomic approaches (Wang et al. 2018; Kalhor et al. 2024) and is likely to improve sensitivity as well as reachable depths. Moreover, 3DEEP is straightforward to integrate with tissue expansion (Chen et al. 2015; Alon et al. 2021). Together with ongoing advances in increasing the capabilities of spatial transcriptomic approaches in characterizing thicker tissue sections (Fang et al. 2023; Gandin et al. 2024 Jan 1; Sui et al. 2024 Jan 1), 3DEEP enables molecular analysis of microscopic cellular structures that, like the hair follicle, are complex along all axes.

Using spatial transcriptomics addressed the key challenges in characterizing hair follicle formation – their asynchronous development and microscopic complexity – as we sampled hundreds of follicles at different stages of development in 3D (**Fig. 2**). Because their transcriptional profile reflected their age, we were able to leverage their asynchronicity from a challenge into an asset by ordering them along their developmental trajectory. This process resulted in a spatiotemporal atlas that animates hair follicle organogenesis from its induction to maturation. It is important to note that our ability to target highly informative genes depended on the wealth of information generated by prior studies (Sennett et al. 2015; Joost et al. 2016; Ge et al. 2020; Joost et al. 2020). While other spatiotemporal atlases have been generated by sampling multiple timepoints in development (Cao et al. 2019; A. Chen et al. 2022; Liu et al. 2022; Sampath Kumar et al. 2023), this work is unique in its temporal resolution, effectively hundreds of time points, which enables one to virtually track structures over time. A similar feat has only been accomplished before without a 3D context and by ordering hundreds of embryos based on morphology prior to dissociation and single-embryo scRNA-seq (Mittnenzweig et al. 2021; Saunders et al. 2023).

Our work captures how the organogenesis phase of the hair follicle results in the emergence of a complex structure and multiple tissues from a simple dermal placode (**Figs. 3 and 4**). While previous histological studies have divided this phase into five stages (Histological Stages 2–6), we conceptualize our animated spatiotemporal atlas here into three Molecular Stages (**Fig. 8**). In the first Molecular Stage, between pseudotimes 0 and 0.2, progenitor cells from all three lineages establish along the main axis of the follicle to form structures and stem cells necessary for formation and long-term maintenance of the follicle. The structures include the dermal condensation at the bottom and uHF at the top of the follicle, which establish its longitudinal axis. The stem cells, which position themselves between the dermal condensation and uHF in the future hair bulge, include keratinocyte stem cells and melanoblasts which migrate from the skin surface. They stratify in heterogeneous layers at this stage (**Fig. 7**). The second Molecular Stage, between pseudotimes 0.2 and 0.7, is marked by differentiation and histogenesis of tissues that undergo destruction and rebirth during the cyclical regenerative life cycle of the hair follicle.

**Figure 8.**
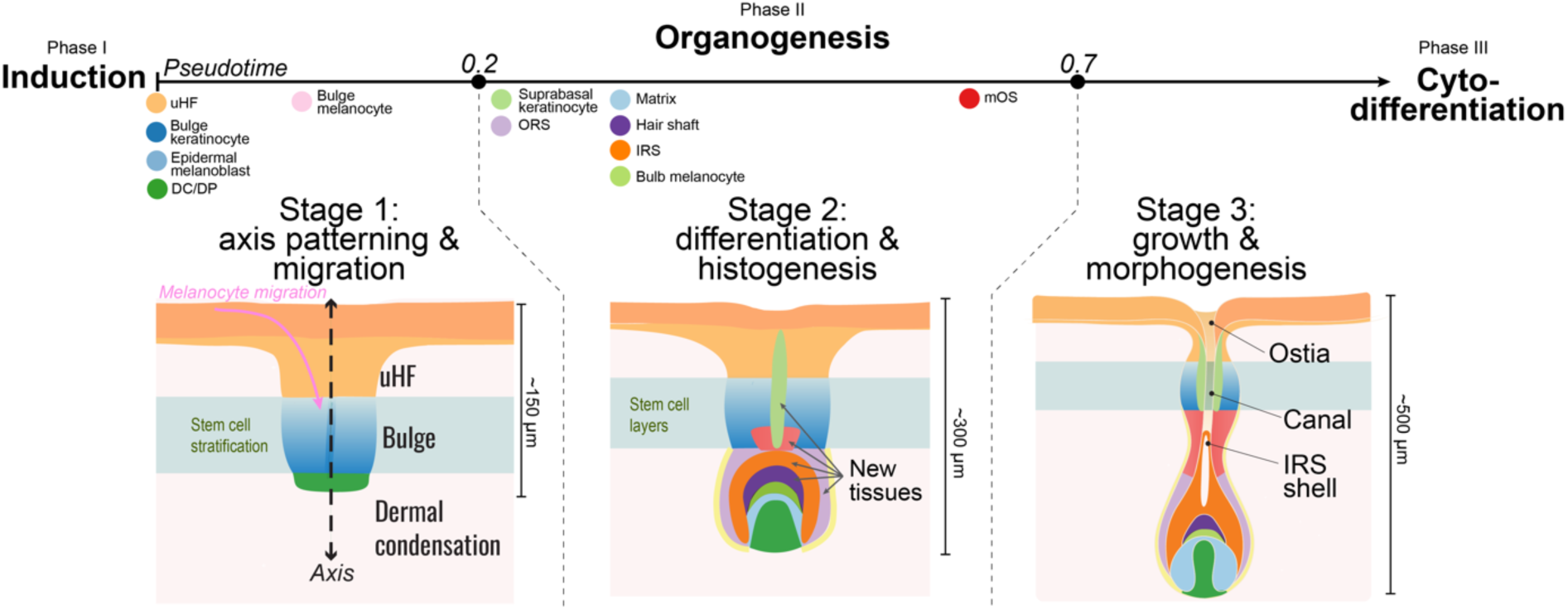
Timeline and molecular stages of hair follicle organogenesis. Illustration summarizing a new model of hair follicle organogenesis in three stages. Hair follicle structure at the end of each of the three molecular stages is shown. Colored circles on the pseudotime axis represent the emergence of each structure.

Several new tissues that are necessary for growing the hair follicle and generating hair strands emerge at 0.2-0.3. These include the DS/reticular fibroblasts, ORS, IRS, hair shaft, matrix, followed by late onset of mOS at 0.6. At this stage, melanocytes are spatially delineated into bulge and bulb melanocytes, settling into their future positions upon the bulb formation. Their orchestrated emergence is a fine example of differentiation and histogenesis in development made possible by germ cells from distinct lineages laying a foundation in an earlier stage. The third Molecular Stage, between pseudotimes 0.7 and 1, is marked by coordinated growth and morphogenesis, when now-formed structures change their morphologies to enable the organ’s main eventual function: hair production. During this stage, a marked increase in the growth of the structure can be detected which is largely driven by the expansion of mOS, ORS, and IRS. As a hallmark of morphological changes, the opening and the canal of the follicle develop through coordinated changes in multiple tissues within the hair follicle (**Fig. 6**).

It is not straightforward to establish the correspondence between the three Molecular Stages we define here and the five Histological Stages defined previously (Schneider et al. 2009) because our results revise the order of events and provide additional information about cellular dynamics that are unattainable without capturing entire follicles. Specifically, we show that ORS emerges first, followed by near-simultaneous inductions of matrix, hair shaft, and IRS. It was previously thought that the matrix emerges first, followed by the gradual acquisition of IRS, ORS, and hair shaft.

Another unique feature of this work is characterizing a large set of morphogen genes in their spatial context during organ formation. Changes in morphogen expression can be captured in scRNA-seq approaches; however, critical information about whether cells in a given location are changing their morphogen expression patterns over time or new cells with a different profile are emerging are lost. The spatial context here allowed us to cluster cell types based on both their transcriptional profile and their position in time and space, which captured morphogen expression changes in structures that are spatiotemporally contiguous (**Fig. 5**). While it is not possible to characterize the function of morphogens in this type of data, two interesting insights emerged from analyzing their spatial distribution. The first insight is the enrichment of uHF and DC/DP, which occupy the bottom of the bulb at the opposite end of the follicle’s long axis, in morphogen expression compared to their surrounding structures. The uHF at the top of the follicle is stable in its morphogen profile whereas DC/DP evolves extensively over time. This observation is interesting because there are more changes during organogenesis in the bulb area of the follicle than there are in the upper parts. Even after the follicle matures, it is known that the upper parts remain steady as the bulb area undergoes destruction and regeneration around the DC/DP during the hair cycle (Schneider et al. 2009). Considering that 1) both DC/DP and uHF emerge very early in hair follicle formation, 2) they are positioned at the extremes of the follicles main axes, 3) they are enriched in morphogens, and 4) their molecular behavior (i.e., stability versus change) presages the behavior of their anatomic neighborhood, we propose that these regions are the main organizers of the hair follicle. This hypothesis expands on the established view of the DC/DP as a main regulator of hair follicle development and growth (Driskell et al. 2011; Morgan 2014). It also suggests the possibility that regeneration of mature hair after its destruction during catagen may involve a sequence of events similar to the second and third Molecular Stages during its initial formation. However, this hypothesis must be explicitly tested in future studies of hair regeneration.

The second interesting pattern that emerged from morphogen analysis is the extensive morphogen shifts in many subpopulations over time (**Fig. 5**). In general, we broadly observed that morphological changes accompanied extensive shifts in morphogen expression profiles. While morphogens have been historically associated with morphological changes, what is striking here is the multiplicity of shifts observed, involving several signaling pathways coordinated in distinct subpopulations. We hypothesize that such shifts are commonplace during the formation of other organs and across development but may be underappreciated due to the low throughput nature of the approaches commonly used to track them.

A few limitations must also be considered when interpreting this spatiotemporal atlas of hair follicle organogenesis. First, these results are based on 81 genes. While these are highly informative genes selected based on the depth of literature in this field and the wealth of scRNA-seq data, it is possible that they do not capture all information and that some tissues or cell types may be missed. Another limitation is that the different subtypes of hair follicles have not been identified here. For example, the first wave of hair follicle induction in mice primarily leads to guard hairs, the second wave primarily to awl hairs, and the third wave to zigzag hairs. While we expect these types to have very similar organogenesis stages because of the shared compartmental structures with few exceptions (Schlake 2005), it is possible that the developmental trajectory presented here could be further deconvolved into multiple follicle subtypes. Another limitation is the lack of detailed lineage information required for understanding the exact origin of tissues that emerge during hair follicle organogenesis.

Addressing this limitation would require incorporating high-throughput lineage barcoding approaches (Kalhor et al. 2018) and is likely to significantly advance our understanding of the cell fate hierarchy during hair follicle formation (Fang et al. 2022). Finally, pseudotime may not have a linear relationship with real time. Therefore, our conclusions about the dynamics of hair follicle growth may be distorted compared to real-time measurements.

Taken as a whole, this study generates an animated atlas of a whole organ’s genesis by integrating time and space using a new spatial transcriptomic approach. This atlas advances our understanding of hair follicle formation and has broad implications for dermatology research. It also deepens our understanding of developmental processes in general because different organs develop based on similar fundamental principles.

## Materials and Methods

### Mouse

C57BL/6J mice (Strain #:000664) were purchased from Jackson Lab. Mice were housed at Johns Hopkins Animal Facility. Animal procedures were approved by the Johns Hopkins University Animal Care and Use Committee (ACUC). Mice were housed in cages on a 12 hour light–dark cycle with accessed water and rodent diet. Animals were monitored by facility staff and researchers. They were euthanized according to institutional guidelines, and death ensured with a secondary means before tissue collection.

### Chemicals and enzymes

All chemicals and enzymes are listed here with the manufacturer and catalog number. DMSO, anhydrous (Biosciences, #786-1666), 10x PBS, pH 7.8 (Bio-Rad, #1610780), 10x PBS, pH 7.4 (Quality Biological, #119-069-131), 1M Tris-HCl, pH 8.0 (Quality Biological, #351-007-101), 20% SDS (Research Products Internatioal, #151-21-3), 1M Tris, pH 7.0 (Invitrogen, #AM9850G), 0.5M EDTA, pH 8.0 (Invitrogen, #AM9260G), Triton X-100 (Sigma-Aldrich, #T8787-100ML), Glycerol (Fisher BioReagents, #BP229-1), Ethanol (Fisher BioReagents, #BP2818-500), Acetic Acid (Sigma-Aldrich, #A6283-500ML), Formamide (Sigma-Aldrich, #47671-250ML-F), Sucrose (Sigma-Aldrich, #S7903-1KG), O.C.T. compound (Tissue-Tek, #4583), Methanol (Sigma-Aldrich, #179337-1L), 20x SSC (Promega, #V4261), VWR Micro Slides (VWR North American, #48300-026), Cover Glass (Corning, #2850-18), Microseal B seal (Bio-Rad, MSB1001), GlassBottom Dish (Cellvis, D60-30-1.5-N), 16% Formaldehyde(Thermo Scientific, #28908), ½ ml Syringe (BD, #305620), PMSF (Cell Signaling Technology, #8553S), DNase I and DNA Digestion Buffer (New England Biolabs, #M0303L), Salmon Sperm DNA (Invitrogen, AM9680), Proteinase K (New England Biolabs, #P8107S), SplintR and SplintR ligase Reaction Buffer (New England Biolabs, #M0375L), Phi29 DNA Polymerase and Phi29 DNA Polymerase Reaction Buffer (New England Biolabs, #M0269L), Thermostable Inorganic Pyrophosphatase (New England Biolabs,, #M0296L), RNase Inhibitor Murine (New England Biolabs, #RNase Inhibitor Murine), Acryloyl-X (Invitrogen, #A20770), Bind Silane (Millipore Sigma, #M6514-25ML), Acrylamide (Bio-Rad, #1610140), Bis-Acrylamide (Bio-Rad, #1610142), N,N,N’,N’-Tetramethylethylenediamine (TEMED, Sigma-Aldrich, #T7024-50ML), Ammonium Persulfate (APS, Sigma-Aldrich, #A3678-100G), Deoxynucleotide (dNTP, New England Biolabs, #N0447S), N,N’-(1,2-Dihydroxyethylene)bisacrylamide (Trolox, Sigma-Aldrich, #294381-5G), Beta-mercaptoethanol (Sigma-Aldrich, #M6250), 5-AA-2’-dUTP (TriLink, N-2049), DAPI (Invitrogen, #D1306). All oligos were ordered from Integrated DNA Technologies.

### 3DEEP-FISH for RNA detection in liver

#### 3DEEP-FISH

Ten-week-old male C57BL/6J mice were sacrificed using CO2. The harvested liver was trimmed to a size of 0.5 cm x 0.5 cm to facilitate PFA diffusion. The trimmed tissue was washed in PBS and incubated in ice-cold 4% formaldehyde/1x PBS, pH 7.4, overnight at 4°C. The fixed samples were washed with PBS (pH 7.4), followed by immersion in 15% sucrose and then 30% sucrose solutions at 4°C until the samples sank in the sucrose solutions. The samples were then embedded in O.C.T. compound, which was solidified using liquid nitrogen. The samples were sectioned at 400 μm using a Cryostat (LEICA CM1860). The slices were transferred to 100% methanol. The methanol-fixed samples were stored at −70°C until use.

Two liver slices were treated with or without DNase I for subsequent 3DEEP-FISH processing. The stored samples were transferred to a 1.5 ml tube containing 1 ml of 2% SSC/8% SDS solution and incubated at 37°C for 30 minutes with a tube rotator. All procedures described in this paragraph were performed with a tube rotator. This wash step was repeated twice. To permeabilize the nuclear membrane, the samples were further incubated in 1 ml of permeabilization solution (2x SSC, 8% SDS) overnight at 37°C. To quench the SDS solution, the samples were incubated in 1 ml of quenching solution (2x SSC, 2% Triton X-100) at 37°C for 1 hour. The solution was then replaced with DNase I buffer (1x DNA digestion buffer, 2% Triton X-100), and the samples were incubated again at 37°C for 1 hour. For the DNase+ control, genomic DNA in the sample was digested by incubating the sample in 1 ml of DNase I solution (1x DNA digestion buffer, 2% Triton X-100, 800 U RNase inhibitor murine, and 400 U DNase I) overnight at 37°C. The DNase-control was incubated in the buffer solution overnight at 37°C. The DNase I solution was quenched with 2x SSC/8% SDS solution at 37°C for 2 hours.

Following gDNA inactivation, we hybridized the RNA in the samples with a padlock oligo. The samples were incubated in 1 ml of hybridization buffer (2x SSC, 20% formamide, 8% SDS, and 20 μg salmon sperm DNA) at 37°C for 1 hour, followed by incubation in the padlock probe hybridization solution (2x SSC, 20% formamide, 8% SDS, 200 nM Apoa2-targeting padlock oligo [/5Phos/AGGTTCATTAAACTGCTGAACATAACAACAAAACAACCTCATTATCTCTCCACACACACTCCTCTCACTCAGGAGCCGGTTTCTCCTCA], 500 nM acrydite-modified RCA primer [/5Acryd/AGTGAGAGGAGTGTGTGTG], and 10 μg salmon sperm DNA) at 37°C for 5 days. The samples were then washed with 1 ml of 1x SSC/0.5% SDS solution at 37°C for 1 hour, twice.

To anchor the hybridized RNA and replace the tissue structure with hydrogel, we initiated the gelation step. A 1 ml monomer solution (10% acrylamide, 0.02% bis-acrylamide, 2x SSC) was prepared. First, the samples were washed with 500 μl of the monomer solution at room temperature for 30 minutes. The remaining monomer solution was bubbled with argon gas to remove oxygen, which inhibits the acrylamide polymerization reaction. To activate the prepared monomer solution, 2 μl of RNase inhibitor murine, 4 μl of 5% TEMED, and 4 μl of 5% APS solutions were added to 200 μl of the argon gas-bubbled monomer solution. The samples and the activated monomer solution were mixed on a glass slide. A coverslip was placed on the samples to form the gel by sandwiching the samples between the glass slide and coverslip. The samples were incubated at 37°C for 1 hour, allowing the gel to solidify sufficiently.

After confirming the formation of a gel, we degraded the tissue structure using Proteinase K. A digestion buffer (50 mM Tris-HCl, pH 7.0, 1 mM EDTA, 2x SSC, 2% SDS) was prepared beforehand. Occasionally, the digestion buffer was prewarmed because 2% SDS can precipitate in the 2x SSC solution. The samples were transferred to the Proteinase K solution (100 μl of Proteinase K, 900 μl of the digestion buffer) and incubated at 37°C overnight with gentle rotation. To quench the Proteinase K and SDS, the samples were washed with 1 ml of quenching solution (2x PMSF, 2x SSC, 0.1% Triton X-100) at room temperature for 30 minutes. The wash step was repeated with 2x SSC/0.1% Triton X-100 at room temperature for 30 minutes, twice, and then with 2x SSC at room temperature for 30 minutes, three times.

Following the gelation and tissue degradation, ligation and RCA steps were performed to amplify oligos using the hybridized padlock probe as a template. First, the samples were briefly washed with ligation buffer (1x SplintR ligase reaction buffer) twice. The samples were then incubated at 37°C for five hours in 500 μl of SplintR reaction solution (1x SplintR ligase reaction buffer, 200 U RNase inhibitor murine, 625 U SplintR) to ligate the padlock probes. Following this, the samples were washed with 1x phi29 DNA Polymerase Reaction Buffer, twice. The samples were incubated in 200 μl of RCA solution (1x phi29 DNA Polymerase Reaction Buffer, 0.1 mg/ml recombinant albumin, 500 μM dNTP, 4 U thermostable inorganic pyrophosphatase, 80 U phi29) at 30°C under agitation.

The amplified concatemers from the RCA reaction were detected with a fluorescent probe (CATAACAACAAAACAACCTCATTATCTCTC/3AlexF647N/). The samples were washed briefly with 2x SSC and equilibrated in 2x SSC/20% formamide. The samples were incubated in a dye probe solution (2x SSC, 20% formamide, 500 nM dye probe, 5 μg/ml DAPI) at room temperature in a dark environment for 2 hours. The samples were then washed with PBS at 37°C for 5 minutes, four times. Finally, the samples were imaged using a Nikon Eclipse Ti2 confocal microscope with a 25X silicone oil-immersion objective (CFI Plan Apochromat Lambda S 25XC, NA:1.05, 599 μm x 599 μm in FOV).

#### Image analysis

The obtained images were denoised using Nikon denoise.ai, followed by deconvolution function within Nikon NIS-Elements. The background signals were subtracted using the default rolling ball background subtraction function in NIS-Elements. The preprocessed images were then used to detect amplicons with the Big-FISH pipeline (Imbert et al. 2022) (https://github.com/fish-quant/big-fish). The signal-to-noise ratio was calculated by dividing the intensities of the amplicons by the standard deviations of disks within 7 to 10 μm of each amplicon.

### Marker gene selection

Candidate marker genes were selected based on a review of relevant literature. Among these candidates, a minimal gene list was curated by confirming their expression and specificity using single-cell RNA sequencing data (GSE131498). Since dataset GSE131498 included E13.5, E16.5, and P0 mouse samples, P0 murine dorsal skin cells (5065 cells) were extracted for this study. To distinguish major skin cell types (keratinocytes, fibroblasts, melanocytes, lymphocytes, adipocytes, and schwann cells), a minimal gene set was selected, ensuring that at least two marker genes were assigned to each cell type. Other cell types, such as lymphatic vessels, were excluded from the analysis due to low marker gene expression levels in the single-cell dataset. For example, *Prox1*, a well-characterized lymphatic vessel marker gene, had fewer than 0.01 reads on average across the cells. As a reference, *Krt14* and *Krt15*, which are widely expressed keratin genes in keratinocytes, had 1.5 and 1.2 reads on average per cell, respectively. We prioritized marker genes within a similar expression range. The literature and databases used for gene curation are summarized in Table 1. Single cells were classified with the candidate marker genes using scSorter (Guo and Li 2021), and the clusters of cell types were visually inspected. To minimize the number of genes used in the study, we manually excluded non-specific or less-abundant marker genes.

After selecting cell type markers, we further curated keratin genes for annotating hair follicle substructures. Joost *et al*. extensively curated keratin gene expressions in these substructures (Joost et al. 2016; Joost et al. 2020). Among the 33 reported keratin genes, 12 keratin genes (*Krt5*, *Krt14*, *Krt15*, *Krt16*, *Krt17*, *Krt25*, *Krt27*, *Krt28*, *Krt32*, *Krt35*, *Krt71*, and *Krt79*) had more than 0.01 reads per cell in the single-cell dataset. These genes are enriched in the outer sheath, IRS, and hair shaft/cortex of hair follicles. We adopted 8 of these keratin genes to detect these major structures. Some keratin genes were excluded due to non-specificity or overlapping expression patterns with other keratin genes. Additionally, a few marker genes were added for their specific expressions (e.g., *Krt6a* in the middle outer sheath) and supplemental structural information (e.g., *Mt1*, expressed in the epidermis, ORS, and matrix, and *Gaba3*, expressed in the epidermis and IRS, related to keratinocyte fate determination).

To broaden the research scope, we included stem cell marker genes and morphogen genes. Stem cell marker genes with evidence of hair regeneration capabilities were curated (Jaks et al. 2008; Lee et al. 2008; Snippert et al. 2010; Brownell et al. 2011). We also included stem cell marker genes used in lineage tracing studies and those with functional relevance to stem cells (Nowak et al. 2008). For morphogen analysis, we focused on the *Wnt*, *Tgf*, *Notch*, *Igf*, and *Edn* families due to genetic evidence supporting their roles in hair development, cycling, and regeneration (Millar et al. 1999; Lin et al. 2000; Reddy et al. 2001; Blanpain et al. 2006; Inoue et al. 2009; Tanimura et al. 2011; Kandyba and Kobielak 2014; Wang et al. 2014; Lim et al. 2016; Rezza et al. 2016; Issa et al. 2017; Trüeb 2018; Bao et al. 2020; Daszczuk et al. 2020; Hochfeld et al. 2021; Li et al. 2022; Simonson et al. 2022; Jacob et al. 2023). Morphogen genes with low expression in the single-cell dataset were excluded from the analysis. Some stem cell-related and morphogen genes were used as markers for keratinocyte/fibroblast substructures (e.g., *Shh* for keratinocyte matrix). These genes were also utilized for cell type/substructure annotations when necessary.

### Padlock probe design

The padlock probe consists of two 20-nt complementary sequences at the 5’ and 3’ ends, which bind to the target RNA sequence. The linker sequence includes a 19 nt RCA primer binding sequence [CACACACACTCCTCTCACT], an 18 nt HybISS bridge probe binding sequence, and a 2 nt gap sequence between the RCA primer binding sequence and the bridge probe binding sequence.

Using a curated list of marker genes, RNA sequences were obtained from the NCBI gene database, specifically using the RefSeq Select RNA sequences. From these sequences, 40 nt candidate RNA sequences were selected by applying a simple threshold: GC content between 40% and 60% for each 20 nt complementary sequence. For the generated 40 nt candidate target sequences, the BLAST command pipeline was used to detect homologous sequences to identify possible non-specific targets. For each suggested non-specific target, the melting temperature was estimated using the Biopython MeltingTemp module. The number of significant non-specific targets was determined by applying a melting temperature threshold of 45°C. The melting temperature of the true positive targets was similarly estimated using the same package.

Given the candidate sequences, GC content, the estimated number of non-specific targets, and the melting temperature of the padlock oligo candidates, we selected favorable probes with 0 non-specific targets, 40% to 60% GC content, and non-repetitive sequences, excluding candidates with CCCC, GGGG, and/or GCGCGCGC motifs. Since we generally obtained more than five probes per gene, we further applied distance clustering to the padlock probe candidates based on their target sequence coordinates on the chromosome. This step was taken to select evenly distributed padlock probe pools, enhancing the chances of probing multiple exons. Five probes were selected for each gene, and these probes were synthesized using the IDT oPools service with 5’ phosphorylation. When fewer than five candidate probes were available, the maximum number of probes was designed for the genes.

The bacterial ribosomal RNA-targeting padlock probes were manually designed. The 16S ribosomal RNA sequence was obtained from the NCBI gene database (NR_024570.1). To probe conserved sequences across bacterial species, each 24-mer was extracted from NR_024570.1 and searched against bacterial species with identical sequences using the BLAST command pipeline. The 16S ribosomal RNA database (Bacteria and Archaea type strains) was used for this analysis. We confirmed 26,810 sequences in the database and calculated the percentages of detected sequences for each 24-mer. The highest detection percentage was 85%, followed by 75%. To increase the melting temperature of the probe bindings, we adjusted the lengths of the probe binding sequences. Since rRNA sequences are also conserved in mammalian rRNA, we checked the mouse rRNA sequences to ensure that the designed probes do not recognize mouse rRNA. Similar to the marker gene padlock probes, the bridge probe binding sequence and RCA primer binding sequence were incorporated. The padlock probes were synthesized using the IDT oPools service.

### HybISS bridge probe design

Unique barcodes for each gene were designed using DNABarcodes (Buschmann and Bystrykh 2013). The original barcodes were generated in the “ACGT” oligonucleotide format, with each 5 nt barcode using 4 channels (ACGT), maintaining a Hamming distance of 2 between each barcode. These barcodes were subsequently converted to a fluorescent format using AF488, AF546, Cy5, and AF750. A total of 144 unique barcodes were generated, with 85 assigned to transcripts and 59 used as negative control barcodes.

To incorporate the fluorescent barcodes into the padlock oligos, we integrated unique bridge oligo binding sequences into the padlock probes for each gene. For this purpose, we utilized bridge probe binding sequences previously designed by Gyllborg and colleagues (Gyllborg et al. 2020). From the 128 pre-designed bridge probe binding sequences available, 85 were randomly selected and incorporated into the respective padlock oligo sequences. These sequences were also used in the corresponding bridge probes to detect the rolonies in the tissue. To ensure the bridge probes would react with the fluorescent probes, the bridge probe sequences were linked to the complementary sequences of the respective fluorescent probes corresponding to the earlier designed barcodes. The bridge probes were synthesized using the IDT oPools service, with the bridge probe sets for each round ordered separately.

### 3D spatial transcriptomics in P0.5 mouse dorsal skin

P0.5 C57BL/6J pups were prepared by crossing male and female C57BL/6J mice, which were harvested between 3 to 6 p.m., defining this time as P0.5. After euthanization, the dorsal skin of the pups was excised with scissors. Fat and muscle tissue were carefully removed using tweezers and a paintbrush, and the samples were then washed with PBS (pH 7.4). The skin was flattened on a filter paper and fixed overnight in ice-cold 4% formaldehyde/1x PBS (pH 7.4) at 4°C. Following fixation, the samples were washed with PBS and transferred to 100% methanol, then stored at −70°C until use.

3DEEP-FISH was applied to the intact skin tissue. First, genomic DNA in the fixed sample was degraded. Stored samples were transferred to 1.5 ml tubes containing 1 ml of 2% SSC/8% SDS solution and incubated at 37°C for 30 minutes on a tube rotator. The wash process was repeated twice. To permeabilize the nuclear membrane, samples were incubated overnight in 10 ml of permeabilization solution (2x SSC, 8% SDS, 1% beta-mercaptoethanol) at 37°C. The samples were then incubated in 2 ml of quench solution (2x SSC, 2% Triton X-100) at 37°C for 1 hour to neutralize the SDS. This was followed by incubation in DNase I buffer (1x DNA digestion buffer, 2% Triton X-100) at 37°C for 1 hour. Genomic DNA was digested by incubating the samples overnight at 37°C in 2 ml of DNase I solution (1x DNA digestion buffer, 2% Triton X-100, 800 U RNase inhibitor murine, and 400 U DNase I). The DNase I solution was quenched by incubating the samples in 2x SSC/8% SDS solution at 37°C for 2 hours.

Following gDNA removal, RNA in the samples was hybridized with a padlock oligo solution. The samples were incubated in 2 ml of hybridization buffer (2x SSC, 20% formamide, 8% SDS, 20 μg salmon sperm DNA) at 37°C for 1 hour. The samples were then transferred to a 96-well plate and incubated in a padlock probe hybridization solution (2x SSC, 20% formamide, 8% SDS, 100 nM padlock oligo each [415 probes, 41.5 μM in total], 100 μM acrydite-modified RCA primer, and 10 μg salmon sperm DNA) at 37°C for 5 overnights with agitation. The plate was sealed with PCR tape to prevent evaporation. After incubation, the samples were transferred to 1.5 ml tubes and washed twice with 1 ml of 1x SSC/0.5% SDS solution at 37°C for 1 hour each.

To anchor the hybridized RNA and replace the tissue structure with hydrogel, the sample was gelled. A 1 ml monomer solution (10% Acrylamide, 0.2% Bis-acrylamide, 2x SSC) was prepared, and samples were washed with 500 μl of the monomer solution at room temperature for 30 minutes. The remaining monomer solution was deoxygenated by bubbling with argon gas to facilitate acrylamide polymerization. The monomer solution was then activated by adding 2 μl RNase inhibitor murine, 4 μl of 5% TEMED, and 4 μl of 5% APS solutions to 200 μl of the argon-degassed monomer solution. The sample and the activated monomer solution were mixed on a slide glass, and a coverslip was placed over the sample to form a gel. The sample was incubated at 37°C for 1 hour to allow the gel to solidify.

Once the gel had solidified, tissue structures were degraded using Proteinase K. Samples were transferred to Proteinase K solution (100 μl Proteinase K + 900 μl digestion buffer (50 mM Tris-HCl pH 7.0, 1 mM EDTA, 2x SSC, 2% SDS)) and incubated overnight at 37°C with a tube rotator. A digestion buffer was occasionally pre-warmed due to SDS precipitation in 2x SSC. To ensure complete digestion, the Proteinase K solution was replaced with fresh solution the next morning, and samples were incubated for an additional 3 hours. Proteinase K and SDS were quenched by washing the samples with 1 ml of quench solution (2x PMSF, 2x SSC, 0.1% Triton X-100) at room temperature for 30 minutes using a tube rotator. The samples were further washed five times with 2x SSC/0.1% Triton X-100 at room temperature for 30 minutes each.

Ligation and RCA were performed to amplify the oligos using the hybridized padlock probes as templates. The samples were briefly washed twice with a ligation buffer (1x SplintR ligase reaction buffer). They were then transferred to a 48-well plate containing 500 μl of SplintR reaction solution (1x SplintR ligase reaction buffer, 200 U RNase inhibitor murine, 625 U SplintR) to ligate the padlock probes. The reaction was carried out under agitation at 37°C overnight. Subsequently, the samples were washed twice with 1x phi29 DNA Polymerase Reaction Buffer and incubated in RCA solution (1x phi29 DNA Polymerase Reaction Buffer, 0.1 mg/ml Recombinant Albumin, 500 μM dNTP, 10 U Thermostable Inorganic Pyrophosphatase, 40 μM 5-AA-2’-dUTP, 160 U Phi29) at 30°C under agitation. The Phi29 solution was replaced with a fresh Phi29 solution 8 hours after the initial reaction, and the RCA reaction was carried out overnight. The total RCA reaction time is approximately 24 hours. The samples were then washed with 2x SSC solution.

Following amplification, the samples were anchored to a glass-bottom plate to minimize deformation during image-based sequencing. A second gelation treatment was performed to anchor the amplicons to a secondary gel and fix the first gel structure on the glass plate. To make the amplicons reactive to the second polyacrylamide gel, samples were treated with 10 mM Acryloyl-X/2x SSC (10% DMSO) at room temperature for 2 hours, modifying the 5-AA-2’-dUTP in the amplicons with acrylamide. A monomer solution (2.5% Acrylamide, 0.125% Bis-acrylamide, 2x SSC, 0.1% Triton X-100) was prepared, with Triton X-100 added after Ar gas bubbling. Samples were washed with 200 μl of the monomer solution. Meanwhile, the glass-bottom plate was activated by sequential washing with water and ethanol, followed by treatment with bind silane solution (1 ml Ethanol, 5 μl Bind Silane, 50 μl Acetic Acid) at room temperature for 30 minutes. The bind silane solution was removed, and the plate was dried. Finally, the plate was washed with ethanol and dried again. The monomer solution was activated by adding 4 μl of 5% TEMED and 4 μl of 5% APS to 200 μl of the monomer solution. The sample was transferred to the activated plate with the activated monomer solution, and a coverslip was placed over it to flatten the sample on the plate. The monomer solution was solidified at 37°C for 1 hour. After solidification, PBS was added, and the coverslip was removed using a ½ ml syringe.

With the fixed sample on the glass-bottom plate, the HybISS protocol was implemented. The sample wash briefly washed with a hybridization buffer (2x SSC, 20% formamide). Bridge probe solutions (2x SSC, 20% formamide, 20 nM bridge probe each [86 probes, 1.72 μM in total], 500 nM AF488 fluorescent probe, 500 nM AF546 fluorescent probe, 500 nM Cy5 fluorescent probe, 500 nM AF750 fluorescent probe) were prepared for five rounds and stored at −70°C until use. The sample was incubated with a bridge probe solution at room temperature for 2 hours in the dark, followed by washing with hybridization buffer (2x SSC, 20% formamide) at room temperature for 10 minutes. The samples were washed six times with PBS at 37°C for 5 minutes each. PBS was then replaced with a scanning solution (1x PBS pH 7.8, 40% (v/v) glycerol, 2 mM Trolox). The sample was imaged for in situ sequencing using a Nikon Eclipse Ti2 confocal microscope with a 20X water-immersion objective (LWD Lambda S 20XC WI, NA: 0.95, 737 μm x 737 μm x 1 μm FOV) equipped with four lasers corresponding to AF488, AF546, Cy5, and AF750 channels. After imaging, bridge probes were stripped using a stripping solution (0.1x SSC, 70% formamide) at 60°C for 10 minutes, 6 times. The hybridization step was repeated for each sequencing round.

### Initial processing of images with ExSeq pipeline

To convert the observed amplicons to transcripts, we first denoised the raw images using Nikon denoise.ai within Nikon NIS-Elements. Background subtraction was performed with the built-in Nikon rolling ball background subtraction function. The processed nd2 files were then converted to the hdf5 file format, optimized for handling 3D array data. Preprocessed images from different cycles were registered using established spatial sequencing analysis pipelines. Image registration and spot detection were conducted using the ExSeq pipeline (https://github.com/dgoodwin208/ExSeqProcessing) (Alon et al. 2021). The coordinates of the amplicons slightly shifted between the imaging cycles were aligned with the ExSeq pipeline. The pipeline parameters were optimized specifically for the images in our study. The color correction feature of the ExSeq pipeline was disabled by setting params.COLOR_CORRECT_CLAMP to 0, as channel shifts in our study were minimal, given that a single camera was used during imaging. Default color correction could introduce artificial channel shifts. Furthermore, we modified the spot detection parameters. The default setting performed spot detection by summing all images from all channels and all rounds. To prevent overcrowding effects caused by image summation, spot detection was performed by using each image from each channel and each round, resulting in a total of 20 images analyzed. Redundant amplicons were eliminated by applying a distance threshold of 10 pixels (3.2 μm) between spots; the same transcripts within 10 pixels were aggregated to prevent double-counting of transcripts.

Using the detected spots, transcripts were decoded, and a confidence score was computed. The ExSeq pipeline provided intensity values for each channel across all rounds at the detected spot coordinates. Scores were calculated using the following equation:

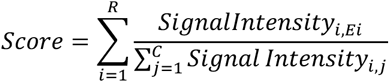

where R is the round, C is the channel, and Ei is the expected code at Round i for a given gene. To evaluate decoding accuracy, we used negative control barcodes that do not correspond to any IDs in the designed probes. Scores were computed for all 85 barcodes, along with 59 negative control codes for each spot. The gene with the highest score was then assigned to each spot.

### Machine learning based sequence decoding

To overcome molecular crowding and spherical aberration, a machine learning-based decoding scheme was applied to our dataset. By applying a score threshold of 4.5 calculated above, a training dataset without false positives was generated. Using the coordinates of the spots with scores greater than 4.5 (top 10% scores), high-quality 8×8 pixel images from the raw sequencing images were extracted. The convolutional neural network pipeline was designed to take 20 images (4 channels x 5 rounds) as input and output one transcript name. Since the number of generated images for each gene was not uniform, the training image data was augmented by altering the order of the image rounds to artificially generate high-quality images for other barcodes. This approach generated 10,000 training data inputs per FOV for 85 barcodes and 59 negative control codes, respectively.

The model was trained in TensorFlow Keras with the generated training dataset, and this process was repeated for six FOVs using the same model. After model training, all amplicon images were processed through the model to predict gene labels. To improve prediction accuracy, the input data was augmented by flipping the images horizontally, vertically, and both ways. These four test image sets were used for prediction, with the trained model returning the likelihood of each gene, and the average likelihood was used to finalize the transcripts.

After decoding, the average negative control barcode percentage was computed for each gene to estimate the fraction of false positives induced by decoding errors. To ensure accurate detection of each gene, the confidence score thresholds were adjusted for each gene so that the average negative control barcode detection rate remained at or below 5% for all genes. Exceptions were *Crabp1*, *Ccl2*, bacterial rRNA, and *Wnt7b*, where the estimated false positive rates induced by decoding errors were 5.7%, 7%, 20%, and 22%, respectively, even at the maximum confidence threshold. These genes were subsequently excluded from the analysis.

### Data processing and analysis for Figures 1J-P

#### Quantification of detected amplicons across skin depth and FOVs

All amplicons detected from six distinct FOVs were used for the quantification. The total number of detected amplicons was quantified for each FOV, and their distribution across varying skin depths was visualized with Matplotlib hist fucntion. Epidermis, dermis, and adipocyte annotations were generated based on the visuzal assessment of the distributions of the keratin genes and adipocyte marker expressions.

#### Histogram analysis of detected amplicons and decoded transcripts

The total number of detected amplicons and the corresponding decoded transcripts were quantified and visualized as a histogram with Matplotlib hist fucntion.

#### Decoding efficiency across skin depth

Decoding efficiency was assessed by calculating the ratio of decoded transcripts to the total number of detected amplicons at each skin depth. The amplicons were divided into 10 bins according to their depth and the decoding efficiencies at each depth were visualized with Matplotlib scatter plot and line plot.

#### Gene distribution analysis across skin depth

The spatial distributions of specific gene categories, including keratin genes, morphogen genes, and stem cell-related genes, were analyzed across skin depths. Detected transcripts across skin depth were smoothed with the statsmodels LOWESS function to illustrate depth-denpendent gene expression for visualization.

#### Correlation between scRNA-seq and 3DEEP-FISH

The detected transcripts with 3DEEP-FISH were compared with a scRNA-seq dataset (GSE131498) to confirm the validity of their expression profile of P0.5 mouse skin. Total numbers of the transcripts for the gene set were plotted with Matplotlib scatter plot. Correlation coefficient and statistical significance were calculated by using the Scipy Spearmanr function.

#### Hair follicle identification and linear transformation

To isolate individual hair follicles from the images, we generated hair follicle mask images to accurately identify the follicle regions. Using Labelme (https://github.com/labelmeai/labelme), an interactive image polygonal manual annotation tool, the contours of each hair follicle were manually marked. Due to the labor-intensive nature of this task, generating masks for every Z-stack is impractical. Therefore, we manually created masks for every 10th Z-stack of the raw *in situ* sequencing images. To enhance boundary detection accuracy for hair follicles, decoded gene coordinates of *Krt6a*, *Krt14*, *Krt15*, *Krt17*, *Krt25*, *Krt27*, *Krt28*, *Krt35*, *Krt71*, *Krt79*, *Mt1*, and *Pmel* were superimposed onto the raw sequencing images. By doing so, we observed the following characteristics: 1. Hair follicles consist of keratin genes at high density rooted in the epidermal layer, 2. Transcript density within hair follicles is higher than in the dermis, and 3. High-density regions exhibit spherical or cylindrical shapes. These observations are consistent with prior knowledge (Joost et al. 2020). After visually confirming these features, all hair follicles, including those partially located at the borders of FOVs, were annotated.

Subsequently, we generated in-between mask labels for every 10th Z-stack by interpolating the manually created masks. This was achieved by transforming the manually generated masks into contours using distance transformation. By connecting these contours between the top and bottom masks with the scipy.interpn function, we computed the intermediate masks. Applying these mask images to the coordinates of the detected spots allowed us to successfully isolate the transcripts from 371 hair follicles across 6 FOVs.

The extracted transcripts from these 371 hair follicles were spatially curved because of their natural shapes. To facilitate structural comparison between follicles, we linearized the hair follicles after excluding partial follicles at the FOV borders. This was accomplished by applying a thin plate spline transformation, converting the original transcript coordinates into a cylindrical shape with a linear Z-axis and a flat XY plane. The thin plate spline method needs “source points” and “target points” to create a warping transformation. We used the centerlines of each hair follicle as source points, as well as two planes representing the skin surface and the bottom of the follicles. The target points were designed as a flat XY plane at Z=0 (for the skin surface), straightened points along the Z-axis (for the hair follicle centerline), and flat XY planes at the corresponding Z coordinates (for the follicle bottom plain). This transformation was then applied to the transcript coordinates to align them along a straight Z-axis with minimal distortion. The detailed steps are as follows:

1. Detect the skin surface using the transcripts from each hair follicle within the bottom 20 μm. The transcripts were segmented into a 5 x 5 pixel XY grid, and the lowest Z value was computed for each grid cell. These lowest values were used to describe the skin surface. This skin surface was subsequently used as a source plane for the thin plate spline transformation.
2. Identify the center point of the skin surface. Since the detected skin surface is tilted, it needs to be flattened to accurately identify its center. To flatten the skin surface, PCA was applied to the points identified in Step 1, successfully producing a flat XY plane. Using this PCA-generated plane, the center point of the skin surface plane was calculated by deriving the mean values of PC1 and PC2. Retrospectively, the corresponding center point in the original data was identified by comparing the PCA plane with the original skin surface.
3. Calculate the angle of the hair follicle and rotate the transcripts to align their overall orientation with the X-axis. To determine this orientation, we used the coordinates of structure-related genes (*Krt6a*, *Krt14*, *Krt15*, *Krt17*, *Krt25*, *Krt27*, *Krt28*, *Krt35*, *Krt71*, *Krt79*, *Mt1*, *and Pmel*). Using the center point and these gene coordinates, the distance of each transcript were computed to select the farthest 100 points. If the total number of transcripts for these genes was fewer than 1000, the farthest one-tenth of the total transcripts were used for the later analysis. We then calculated the angles from the center point to the farthest points. The average angle estimated from the 100 farthest points was applied to rotate the transcripts onto the X-axis. Additionally, the average coordinates of the farthest points were used later as an endpoint for the centerline calculation.
4. Estimate the centerline using the coordinates of the transcripts in both the XZ and XY planes, which were later combined to construct a 3D centerline. First, from the transcript coordinates, we generated a binary mask array describing the hair follicle and non-follicle regions. This conversion utilized Alphashape, which outlines a boundary area from 2D points. From the resulting binary image, hair follicle contours were identified using the skimage.find_contours function. The centerline was then estimated from these contours using the Voronoi diagram. The Voronoi diagram subdivides a 2D area into multiple regions, ensuring each region corresponds to the closest data point. This method effectively identified central points within the contours of the hair follicle. To improve the accuracy of the centerline inference, these central points were computed for all transcripts and selected transcripts (*Krt6a*, *Krt14*, *Krt15*, *Krt17*, *Krt25*, *Krt27*, *Krt28*, *Krt35*, *Krt71*, *Krt79*, *Mt1*, *and Pmel*), respectively. These separately calculated central points were combined to improve the inference of the centerline. Using the points around the center of the hair follicle, a centerline was derived by computing its shortest path from the start point to the endpoint obtained in Steps 2 and 3. The derived path was smoothed using the statsmodels.lowess function.
5. Detect the vertical plane at the bottom of the hair follicle. With the centerlines of the XZ and XY planes acquired, the 3D centerline can be described as a function of X. Using the endpoint (X_end, Y_end, and Z_end) of the centerline and the second point (X_end - 40 μm, Y at (X_end - 40 μm), and Z at (X_end - 40 μm)), the vertical plane at the hair follicle bottom was computed using a vector equation of a plane. The generated vertical plane at the follicle bottom was used as the source points. When the overall length of the hair follicle in the X coordinate was less than 40 μm, half of the maximum value of X was used to generate the vertical plane at the hair follicle bottom.
6. Obtain the target plane and adjust the target plane angle. With the surface plane, centerline, and hair follicle-bottom vertical plane prepared, we next produced the corresponding target planes with adjusted angles. Applying PCA to the source planes identified in Steps 1 and 5 produced flat planes, which served as the target planes for the skin surface and the bottom vertical planes, respectively. Since the angles of the PCA planes were arbitrary, the target planes were rotated so that the transcripts in the direction of the hair follicles align with the X-axis. To achieve this, the direction of the hair follicle was calculated using the centerline on the XY axis. With the given angle, the target plane was rotated.
7. Prepare the summarized source points and target points. The source points were generated in Steps 1, 4, and 5. For the source point planes, the target points were generated in Step 6. For the remaining centerline, the target points were generated using a simple numerical derivative. Here, the target points of the centerline are equivalent to the linearized centerline. We calculated the length of the centerline with a step size of 0.1 in X using the following equation:

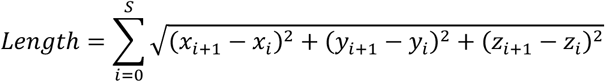

S is the total number of steps. For each step, the corresponding target points were generated on the linear Z axis with the equivalent lengths. For the skin surface target plane, Z was set at 0, while for the vertical plane at the hair follicle bottom, the respective Z coordinate was used, equivalent to the overall hair follicle length.
8. Transform the coordinates of the transcripts. By using the source points and target points, a thin plate spline warping function was trained. Applying this function to the coordinates, the transformed coordinates of the transcripts around the Z-axis were computed.

### Data processing and analysis for Figures 2B-N

#### Hair follicle density analysis

The distribution of full and partial hair follicles was visualized using the Matplotlib scatter plot function. To avoid double-counting due to partial hair follicles that span across multiple FOVs, 15% of the data from the sides was excluded from the density estimation. Focusing on the hair follicles located at the center of the FOVs, the density was calculated by dividing the number of hair follicles by the area across six FOVs.

#### Transcript quantification per hair follicle

After assigning the transcripts to each hair follicle with the masks, the number of transcripts detected within each hair follicle was visualized with the Matplotlib hist function.

#### Hair follicle length calculation and distribution

After linearizing the hair follicles, their lengths were calculated by subtracting the minimum axial coordinate Z of the transcript from the maximum axial coordinate Z. The distribution of these lengths was then plotted using the Matplotlib hist function.

#### Hair follicle pseudotime inference

To evaluate the developmental trajectory of the identified hair follicles, we applied single-cell RNA sequencing analysis techniques using the Seurat and Slingshot pipelines (Satija et al. 2015; Street et al. 2018). Using the mask images created in the previous section (“Hair follicle identification and linear transformation”), the transcripts which belong to each hair follicle were identified. This transcripts-hair follicle relationships were converted to a count table. Four genes (*Crabp1*, *Ccl2*, Bacteria, and *Wnt7b*) were excluded because they did not meet the confidence threshold criteria, which was set to allow a maximum false positive rate of 5%. The count matrix was normalized and scaled using the scTransform function in Seurat (Hafemeister and Satija 2019). Using the normalized count matrix, 10 principal components were computed. The dimension reduction plot visually showed one developmental trajectory. In our study, all hair follicles were artificially assigned to a single cluster using the FindClusters function in Seurat since they are categorically the same organ. The Slingshot pipeline was then used to estimate the developmental trajectory within the cluster. The pseudotime was obtained using the slingPseudotime function.

#### Correlation analysis between pseudotime and hair follicle lengths

The correlation between pseudotime and hair follicle lengths was examined with the Scipy Spearman function. The correlation was plotted with the Matplotib scatter plot.

#### Secondary-tertiary classification of hair follicles

The hair follicles were classified into secondary and tertiary categories based on the Gaussian mixture model clustering on pseudotime. Pseudotime distribution showed the bimodal peaks. The cluster with higher pseudtime was annotated as secondary hair follicle whereas the cluster with lower pseudotime was termed tertiary hair follicle.

#### Correlation analysis between pseudotime of neighboring hair follicle pairs

By using the Scipy KDTree function, the nearest tertiary hair follicles of the secondary hair follicles within 100 μm range were determined. Pseudotime correlation of these secondary-tertiary hair follicle pairs were examined with the Scipy Pearsonr function. The linear fitting curve between pseudotime of these pairs was obtained with the Scipy ODR function.

#### Squared error analysis

Linear fitting curves between pseudotime and hair follicle lengths for secondary, tertiary, and all hair follicles were obtained with the Sklearn LinearRegression function. With the given linear curves, squared errors between the predicted pseudotime from the lengths and the observed pseudotime were computed by using the following equation:

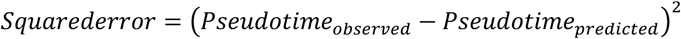

The calculated squared errors for the three groups were compared using a two-sided Wilcoxon rank sum test.

#### Correlation analysis between gene expressions and pseudotime

Relationships between normalized expression levels of genes in hair follicles and psuedotime of hair follicles were examined. First, the transcript counts in hair follicles were normalized with the Seurat scTransform function. Then, correlations between the normalize expressions and pseudotime were calculated for each gene with the Scipy Pearsonr function. The normalized expressions of the top 40 genes with the lowest p-values were visualized with the Matplotlib imshow function.

### Data processing and analysis for Figures 3A-N

#### Baysor single cell segmentation

To establish single-cell level analysis, Baysor was applied to the decoded transcripts to infer cell boundaries. Baysor requires the coordinates, gene names, expected radius of the cells, standard deviation of the radius, minimum molecules per cell, and the number of cell type clusters. The XY coordinates of the transcripts in pixels were scaled to μm before Baysor. The parameters used in the study were as follows: radius at 5 μm, standard deviation of the radius at 50%, minimum molecules per cell at 3, and the number of cell type clusters at 9. The parameter was obtained by measuring the radius of spatially isolated dermal fibroblasts in our dataset. The cellular radius observed in our study ranged between 5 to 10 micrometers.

#### Distribution analysis of the number of inferred cells per hair follicle

The number of inferred cells within each hair follicle was quantified and visualized with the Matplotlib hist function.

#### Correlation analysis between hair follicle lengths and the number of inferred cells

The correlation between hair follicle lengths and the number of inferred cell using Baysor was plotted with the Matplotib scatter plot.

#### Distribution analysis of the number of transcripts per inferred cell

The number of transcripts per inferred cell was quantified and visualized with the Matplotlib hist function.

#### Sensitivity comparison analysis between 3DEEP-FISH and UMI-based scRNA-seq

Gene expression levels per cell was compared between the 3DEEP-FISH dataset and publicly available UMI-based scRNA-seq data (GSE108097). For this comparison, mean gene expressions per cell for a shared set of genes was calculated for both datasets. A box plot was used to visualize the comparison, and the statistical significance of differences in methodological sensitivity was evaluated using a Wilcoxon signed-rank test.

#### Cell type annotation of hair follicles

Using the curated marker genes, the inferred cells were classified into fibroblasts, keratinocytes, and melanocytes based on their transcriptomes. Cells with fewer than three transcripts were excluded from the analysis. The count table of transcripts per cell was log-transformed and subsequently z-score transformed. Using the transformed count table, scSorter separated the cells into fibroblasts, keratinocytes + melanocytes, and unknowns (Guo and Li 2021). Unknowns were excluded from further analysis. Among keratinocyte and melanocyte cells, those cells positive for melanocyte markers (Pmel, Dct, and Ptgds) with ≥2 transcripts of any marker gene were annotated as melanocytes. This approach was taken due to the high abundance of keratin genes mixed with melanocyte marker genes and the inherent 5% false positive rate. The validity of this approach was confirmed by examining the spatial distribution of melanocytes shown in Figure 7A, which is consistent with previous reports (J. Chen et al. 2022).

#### Structual annotation of hair follicles

Using the detailed spatial distributions of marker genes previously reported (Rezza et al. 2016; Joost et al. 2020), the structures of developing hair follicles were manually annotated because of the limited number of genes available. First, keratinocyte cells were classified into IRS, hair shaft, matrix, and outer sheath keratinocytes using density-based spatial clustering (DBSCAN) applied to marker gene-positive cells, curated based on their spatial information summarized in Table 1. Specifically, 12 genes were used for this classification: *Krt35* for hair shaft; *Krt25*, *Krt27*, *Krt28*, and *Krt71* for IRS; *Shh* and *Mt1* for matrix; *Krt79* for uHF; *Barx2* and *Krt14* for uHF-mORS; *Krt17* for uHF-ORS; and *Krt6a* for mORS. DBSCAN was applied to the 3D coordinates (axial position Z, radial position R, and rank of hair follicle pseudotime) of the marker gene-positive cells. High-density regions of the cells were identified as the structures denoted above, including IRS, hair shaft, and matrix. *Krt79*, *Barx2*, *Krt14*, *Krt17*, and *Krt6a+* cells spatially overlapped, as they cover parts of the outer sheath keratinocytes, and thus, they were summarized as outer sheath keratinocytes. When the same cells were annotated with different labels, the quantiles of the expression levels were compared, and the structures of the marker genes with relatively high quantiles were assigned to these cells. After the density-based annotations, some keratinocyte cells, which were not assigned to any structures, had widely expressed keratin genes, such as *Krt15*, without the expressions of structure-related marker genes. These keratinocyte cells were assigned to outer sheath keratinocytes if their nearest cells are outer sheath keratinocytes. These neaerest cells were evaluated by using the SciPy KDTree function.

Following the density-based clustering, the outer sheath keratinocytes were further classified into uHF, bulge keratinocytes, suprabasal keratinocytes, mOS, and ORS. *Krt79*, *Barx2*, *Krt14*, *Krt17*, and *Krt6a*+ cells spatially overlapped, displaying spectra of keratin genes featured by the addition and also subtraction of certain genes. Therefore, a transcriptome-based approach was applied. Given that the average number of transcripts per cell in our dataset is ∼16, and a conservative radius parameter was used for Baysor single-cell inference, single-cell transcriptome clustering could be biased, possibly producing artifical clusters. To mitigate this issue and interpret the spatial representation of hair follicle structures, we generated pseudobulks from the outer sheath keratinocytes by applying K-means clustering to the 4D coordinates (axial position Z, X, Y, and hair follicle pseudotime) of the outer sheath keratinocytes. From these pseudobulks, a transcript count table was generated. Among the geneset, Fibroblast markers *Col6a1*, *Col6a2*, *Trps1*, and *Fbln2* were excluded from the analysis to prevent confounding effects from neighboring fibroblast populations. The count table was transformed using scTransform in Seurat, followed by Leiden clustering (Traag et al. 2019). The clustering results were visualized using UMAP (McInnes et al. 2020). Based on their spatial distributions and enriched marker genes, the clusters were termed uHF, bulge keratinocytes, suprabasal keratinocytes, mOS, and ORS.

Similarly, the fibroblasts were also classified into papillary/IF fibroblast, bulge fibroblast, DS/Reticular fibroblast, and DC/DP. Fibroblast pseudobulks were generated by applying K-means clustering to the 4D coordinates (axial position Z, X, Y, and hair follicle pseudotime) of the fibroblasts. A transcript count table was generated after removing keratinocyte marker genes (*Krt14*, *Krt17*, *Krt15*, *Mt1*, *Gata3*, *Barx2*, *Krt79*, *Krt28*, *Krt35*, *Krt27*, *Krt71*, *Krt25*, *Krt6a*, *and Col17a1*) to prevent confounding effects from neighboring keratinocyte populations. The count table was transformed using scTransform in Seurat, followed by Leiden clustering. The clustering results were visualized using UMAP. Based on their spatial distributions, the clusters were termed papillary/IF fibroblast, bulge fibroblast, DS/Reticular fibroblast, and DC/DP.

Lastly, the melanocytes were classified into epidermis melanoblast, bulge melanoblast, and bulb melanoblast. Melanocyte pseudobulks were generated by applying K-means clustering to the 4D coordinates (axial position Z, X, Y, and hair follicle pseudotime) of the melanocytes. A transcript count table was generated without keratinocyte and fibroblast markers. The count table was transformed using scTransform in Seurat, followed by Leiden clustering. The clustering results were visualized using UMAP. Based on their spatial distributions, the clusters were termed epidermis melanoblast, bulge melanoblast, and bulb melanoblast.

### Data processing and analysis for Figures 4A-P

#### Cellular population dynamics analysis in hair follicle development

Cell number changes over pseudotime were visualized to illustrate the overall dynamics of cellular proliferation and migration. The number of cells per hair follicle in different structures over pseudotime was visualized using a Matplotlib scatter plot. The statsmodels LOWESS function was applied to the scatter data points to smooth the population changes over pseudotime. A stream graph was generated based on the LOWESS results for each structure, with the width of the stream reflecting the number of cells. For better visualization, the width for melanocyte lineage was slightly adjusted, because melanocyte populations were relatively small and their stream changes could be difficult to see.

### Data processing and analysis for Figures 5A-F

#### Heatmap visualization of gene enrichment and expression levels

The cells within each structure were divided into pseudotime bins of 0.1 intervals. Because there are a limited number of cells before 0.2, the cells with pseudotime lower than 0.2 were categorized as one group. Similarly, for the structures which are derived in the middle of hair follicle development, the cells with pseudotime lower than 0.5 were classified as a group. For each group of cells, highly expressed genes were identified using permutation tests. For 40 selected morphogen and signaling molecule genes, the mean expression level in each grouped cell set was compared with the mean expression level in all cells. If the expression level in the group was higher than the overall average, permutation test was applied. Here, a random sample of cells, which is equal in number to the grouped cells, was drawn from the entire dataset, and the mean expression level was recalculated. This sampling process was repeated 100,000 times to compute statistical significance. Specifically, the observed mean expression in the group and its corresponding quantile among the sampled values were used to calculate the p-value. The computed p-values were then adjusted to q-values using the Benjamini-Hochberg procedure. To visualize the wide range of q-values, log10(1/q-value) was used, with clipping set to a maximum of 4. Expression levels were log-transformed to account for the wide variance across genes and then converted to z-scores. Z-score values were clipped at 1 to enhance interpretability and emphasize significant expression changes.

#### Quantification of enriched genes across structures

To capture dynamic and stable nature of morphogen gene regulations in structures, the number of enriched genes were counted for each structure. Genes with a q-value less than 0.05 in at least one pseudotime bin as well as in all pseudotime bins were separately counted. The Matplotlib scatter plot function was used to visualize the number of enriched genes within each annotated structure. The expression levels were log-transformed, followed by z-score transformation.

#### Variance analysis of enriched morphogen gene regulations across pseudotime

The enriched morphogen genes for each structure were selected based on a criterion: a q-value of less than 0.05 in at least one pseudotime bin. The variance of morphogen expression levels was calculated as follows: 1) transcript counts were log-transformed, followed by z-score transformation; 2) for each time bin, the mean z-score was calculated; and 3) the variance of mean z-scores across pseudotime bins was determined. This process was repeated for all enriched genes in each structure. The Matplotlib boxplot function was used to visualize the calculated variance. Statistical significance between uHF and DC/DP was assessed using a two-sided Wilcoxon rank-sum test. The mean variance calculated for each structure was used in Figure 5D. The structures were grouped into upper and lower parts. The upper structures include uHF, suprabasal, bulge keratinocyte, papillary/IF fibroblast, and bulge fibroblast, while the lower structures include ORS, IRS, hair shaft, matrix, DS/reticular fibroblast, and DC/DP. The mean variance of the two groups was visualized with the Matplotlib boxplot function. Statistical significance between the upper and lower parts was assessed using a two-sided Wilcoxon rank-sum test.

#### Density maps of annotated structures

The cell densities of the annotated structural cells across pseudotime and axial position Z were visualized using the Matplotlib 2D histogram function. The boundary lines of the annotated structures were determined based on the coordinates of the annotated cells. Specifically, for each pseudotime value, the cells in the top and bottom 5% of axial positions were identified. A LOWESS fitting curve was then applied to these cells to estimate the boundary lines of the structures.

#### Correlation analysis between gene expression levels and the number of cells over pseudotime

To explore the relationship between gene expression and cellular population dynamics, we calculated correlation coefficients between the number of cells and the mean expression levels over pseudotime. The cells were divided into 50 equal-sized bins based on their pseudotime, and the mean expression levels per cell were examined within each bin. These mean expression levels were then compared to the number of cells at the corresponding pseudotime. Correlation coefficients were computed using the Scipy pearsonr function. Genes with correlation coefficients greater than 0.5 or less than −0.5 were visualized using the Matplotlib imshow function.

### Data processing and analysis for Figures 6A-I

#### Classification of hair follicles by ostia formation

To classify ostia formation without computational bias, visual assessments were used since computational analysis based on trends over pseudotime could overlook the individual shapes of hair follicles. Eight representative images of hair follicles were selected to illustrate the formation of hair follicle ostia in Figure 6A, illustrating the morphology of hair follicles on which their ostia were visually assessed. To evaluate the diameter of growing ostia, the distribution of *Krt79*+ cells (≥ 2 *Krt79* transcripts) was analyzed along the radial axis R below the hair follicle surface (within 100 μm from the surface) and across pseudotime. A scatter plot was used to visualize the radial distribution of these cells in relation to pseudotime, with the ostia boundary estimated using LOWESS fitting. To model the trend of the smallest R values over time, cells were grouped by time points, and within each group, the mean of the five smallest R values was calculated. These values were then smoothed using the LOWESS function. The LOWESS output was rescaled by normalizing its range between the minimum and maximum values to standardize the curve for visualization.

#### Visualization of cellular distributions of IRS, hair shaft, and outer sheath keratinocyte in pseudotime bins

The distributions of IRS, hair shaft, and outer sheath keratinocyte cells were examined across pseudotime bins along both axial and radial coordinates. Cells were binned based on their pseudotime, and scatter plots were generated to visualize their spatial distributions within the hair follicles.

#### Classification of hair follicles by IRS cylindrical shell formation

To classify IRS cylindrical shell formation without computational bias, visual assessments were conducted. Due to the smaller radius of the IRS cylindrical shell compared to the ostia radius, computational approaches are less effective for this analysis. Eight representative images of hair follicles were selected to illustrate the formation of the IRS cylindrical shell in Figure 6E, highlighting the morphology of these structures, which are characterized by green amplicons in the images.

#### Analysis of structural positions over pseudotime

Since ostia and canal formation appeared to be related to IRS movement, the movements of these structures in relation to others were visualized. The spatial distributions of the annotated structural cells across pseudotime and axial position Z were displayed. To illustrate the boundary lines of the annotated structures, the coordinates corresponding to the top and bottom 5% of axial position values of the annotated cells were used. A LOWESS fitting curve was then applied to these coordinates to estimate the boundary lines of the structures.

#### Assessment of length of the structures over pseudotime

The lengths of each structure were determined by subtracting the minimum value from the maximum value of the axial coordinate Z for each structure within each hair follicle. The mean lengths of the hair follicle structures were then calculated for each pseudotime bin. A Matplotlib scatter plot was used to illustrate the temporal changes in these lengths.

#### Analysis on gene expression changes associated with ostia and IRS shell formation

The mean expression levels per cell for each hair follicle were evaluated for genes showing temporal changes between pseudotime 0.5 and 1. The mean expression levels per cell for each hair follicle were visualized using box plots. Statistical significance between the time points was assessed with a two-sided Wilcoxon rank-sum test.

#### Gene expression analysis associated with ostia and IRS cylindrical shell formation

To analyze gene expression level changes associated with ostia and IRS cylindrical shell formation, genes showing temporal variation were selected. The expression levels of morphogen genes were compared between adjacent temporal bins (0.5-0.55, 0.55-0.6, 0.65-0.70, 0.70-0.75, 0.75-0.80, 0.85-0.90, 0.95-1.0) using a two-sided Wilcoxon rank-sum test. The four most significant genes were visualized using Matplotlib’s box plot function.

### Data processing and analysis for Figures 7A-K

#### Tracking melanocyte cell positions over pseudotime

The axial positions of melanocyte cells were tracked over pseudotime using the Matplotlib scatter plot function. A scatter plot was created to visualize the distribution of melanocyte cells along the axial axis across pseudotime, with three subtype annotations: epidermis melanocytes, bulge melanocytes, and bulb melanocytes.

#### Distributions of stem cell marker-positive keratinocytes over pseudotime

The axial positions of keratinocytes expressing stem cell markers (*Cd34*, *Gli1*, *Lgr5*, *Lgr6*, and *Lrig1*, with ≥2 transcripts per cell) over pseudotime were visualized using the Matplotlib scatter plot function. The color of each data point was adjusted based on the spatial density value of each cell, estimated with Scipy’s gaussian_kde function. Kernel density estimation was applied to the 2D coordinates of the axial position Z and the rank of hair follicle pseudotime to infer the density values at the given positions.

#### Estimation of representative stem cell positions using local regression

To determine the representative positions of stem keratinocytes in hair follicles over pseudotime, the highest density regions of stem cell marker-positive cells were first identified using the density values estimated above. Stem cells with the highest density in each hair follicle were extracted. The spatiotemporal positions of these cells were then smoothed using the statsmodels LOWESS function, providing the representative positional trajectories of stem marker-positive cells.

#### Cellular population dynamics analysis of keratinocyte stem cells

The number of stem cell marker-positive keratinocytes per hair follicle for each stem cell marker (*Cd34*, *Gli1*, *Lgr5*, *Lgr6*, and *Lrig1*, with ≥2 transcripts per cell) over pseudotime was visualized using a Matplotlib scatter plot. The statsmodels LOWESS function was applied to the scatter data points to smooth the population changes over pseudotime.

#### Distribution of *Col17a1*+ keratinocytes over pseudotime

To visualize the distribution of *Col17a1+* keratinocytes (≥2 transcripts per cell) over pseudotime, the spatiotemporal coordinates of these cells within the hair follicles were plotted using the Matplotlib scatter plot function. The axial positions Z and pseudotime were used as coordinates. Kernel density estimation was applied to the 2D coordinates of axial position Z and the rank of hair follicle pseudotime to infer density values at the given positions. The estimated densities were then superimposed onto the scatter plot data points.

#### Differential expression analysis between keratinocyte stem cells and non-stem cells in the bulge

To examine the expression profiles of bulge stem cells compared to bulge niche keratinocytes, all keratinocytes located in the bulge were first identified. Using the coordinates of the bulge keratinocytes identified in our study, the boundary lines of the bulge region were determined. Specifically, for each pseudotime value, the bulge keratinocytes in the top and bottom 5% of axial positions were identified. A LOWESS fitting curve was applied to these cells to estimate the boundary lines of the structures. All keratinocytes within these boundary lines were then extracted for further analysis. Among the extracted cells, stem cell marker-positive keratinocytes (*Cd34*, *Gli1*, *Lgr5*, *Lgr6*, and *Lrig1*, with ≥2 transcripts per cell) were classified as stem bulge keratinocytes, while the remaining cells were classified as niche bulge keratinocytes. Differentially expressed genes between these groups were identified using a two-sided Wilcoxon rank sum test. These p-values were then adjusted to q-values using the Benjamini-Hochberg procedure. For visualization, the q-values and log2(fold change) were plotted by using the Matplotlib scatter function. The q-values were clipped at 3 for consistent visualization.

### Data processing and analysis for Figures S1-3

#### Visualization of morphogen gene density distributions

To analyze the spatial distribution of gene expression over pseudotime and the axial coordinate Z, we first extracted the 2D coordinates (pseudotime and axial coordinate Z) of the corresponding transcripts for all hair follicles and each structure. Since the distribution of pseudotime is not uniform, the estimated density is biased. Therefore, pseudotime was transformed to ranks. Using the pseudotime ranks, the axial coordinate Z, and the np.histogram2d function, density maps of morphogen genes were generated. The values in the generated density maps were then scaled to the original pseudotime. These scaled values were plotted using the Matplotlib plt.pcolormesh function.

## Acknowledgments

We thank Dr. WanYing Lin, Jinwoo Jun, and Caroline Ghio for their feedback and suggestions throughout the course of the project. We thank Drs. Kian Kalhor, Jean Fan, Justus Kebschull for their input on the project and critical reading of the manuscript. We also thank Dr. Daniel R. Goodwin for assisting in the adoption of *in situ* sequencing decoding pipelines, Dr. Sashank K. Reddy for helpful discussions, and Dr. Andrew Ewald for the use of a 20X water-immersion objective. We also thank Rachel Boyd for diffusion time modeling.

## Author contributions

S.A., L.A.G., and R.K. conceptualized the study, S.A. designed and performed experiments, S.A. and C.Y. analyzed the data, R.K. and S.A. wrote the manuscript with feedback from L.A.G., L.A.G. and R.K. supervised the study.

## Data and materials availability

All pseudotime, coordinate, processed gene expression, and decoded *in situ* sequencing data and python and R scripts to reproduce the results are available upon request from the authors.

## Supplementary Figures

**Figure S1. Related to Figure 5: Characterizing morphogen gene expression over space and time.** (**A**) Density maps of keratin and morphogen genes across pseudotime and arong the axial coordinate Z.

**Figure S2. Related to Figure 5: Characterizing morphogen gene expression over space and time.** (**A-H**) Density maps of morphogen genes in keratinocyte structures, including uHF, bulge keratinocyte, suprabasal keratinocyte, mOS, ORS, IRS, hair shaft, and matrix, across pseudotime and arong the axial coordinate Z.

**Figure S3. Related to Figure 5: Characterizing morphogen gene expression over space and time.** (**A-D**) Density maps of morphogen genes in fibroblast structures, including papillary/IF fibroblast, bulge fibroblast, DS/Reticular fibroblast, and DC/DP, across pseudotime and arong the axial coordinate Z.

## Supplementary Tables

**Table S1. List of padlock oligo sequences and thermodynamic characteristics.** The file separated submitted as an auxiliary supplementary material because of the data size.

**Table S2. List of bridge probes and assigned ID sequences.** The file separated submitted as an auxiliary supplementary material because of the data size.

**Table S3. Summary of hair follicle positions.** The file separated submitted as an auxiliary supplementary material because of the data size.

