## Supplementary figures and images for "Deconvolving organogenesis in space and time via spatial transcriptomics in thick tissues"

### Figure S1

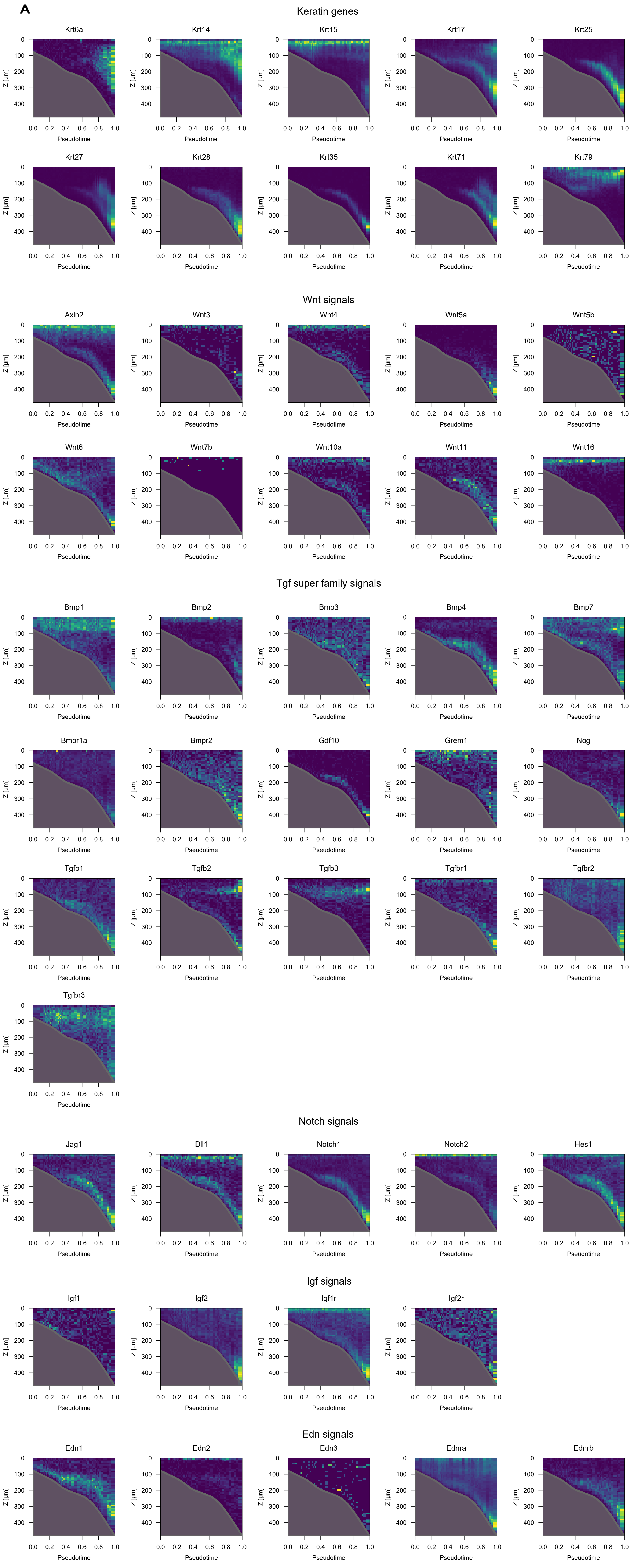

### Figure S2

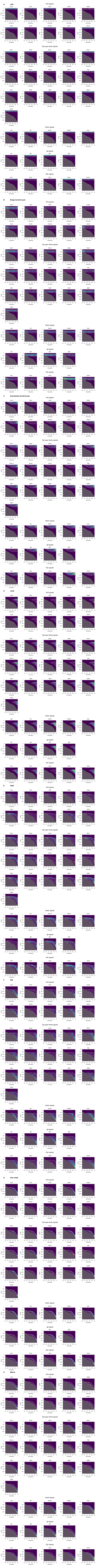

### Figure S3

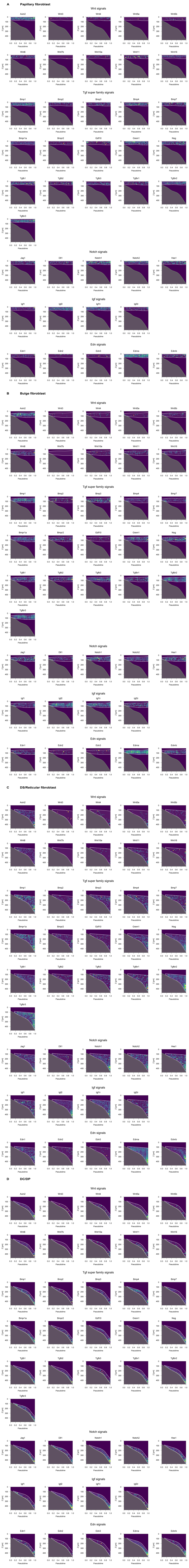
